# The mechanics and physics of tofu: understanding hydrated soft solids through feature networks

**DOI:** 10.64898/2025.12.10.693552

**Authors:** Birte Boes, Jaan-Willem Simon, Hagen Holthusen, Ellen Kuhl

## Abstract

Tofu remains one of the world’s most important plant-based foods, famous for its cultural legacy, nutritional benefits, and environmental sustainability. Made up of only two ingredient–soy beans and water–it has nourished people for centuries; yet, its rheology remains incompletely understood. Here, we use tofu as a minimal model system to study how composition governs the mechanics and physics of hydrated soft solids. We perform over a hundred controlled compression tests on silken, firm, and extrafirm tofu, and observe a strong nonlinearity, a pronounced viscoelasticy, and a more than ten-fold increase in stress with only six percent decrease in water content. We demonstrate that automated model discovery can identify inelastic constitutive laws that accurately capture these unique characteristics across multiple loading magnitudes and rates. The discovered models autonomously separate elastic and inelastic mechanisms and reveal that tofu elasticity depends primarily on the second isochoric invariant, while tofu inelasticity depends on a combination of the second and third deviatoric invariants. This functional form is universal across all three tofu types, but scales in magnitude with water content. Our new water-content-based feature network reveals that this correlation in not linear, as assumed by traditional bi-phasic theories, but rather highly non-linear and sensitive to the individual model terms. These results position tofu as a quantitative benchmark for nonlinear poroviscoelastic solids and demonstrate how physics-informed machine learning can uncover the constitutive structure of hydrated soft materials. Our source code and examples are available at https://doi.org/10.5281/zenodo.16993236.

## 1. Introduction

Tofu feeds more than a billion people worldwide and has done so for over two thousand years [68]. It is one of the simplest and most influential plant-based foods in human history [80]. Beyond its cultural role, tofu promotes human health [63], contributes to planetary sustainability [62], and remains the cornerstone of a modern plant-based diet [66]. Yet, the material itself is remarkably simple [41]: Soymilk and a mineral coagulant form a hydrated protein gel with only two essential ingredients, soybeans and water [84]. Whether we make our own tofu at home or buy it in supermarkets, Figure 1 illustrates that its texture can range from silken to extrafirm, determined almost entirely by its water content [13] and by the specifics of its coagulation process [81]. This simplicity makes tofu an ideal prototype material for testing, validating, and revealing the power of automated model-discovery for soft matter systems.

**Figure 1.**
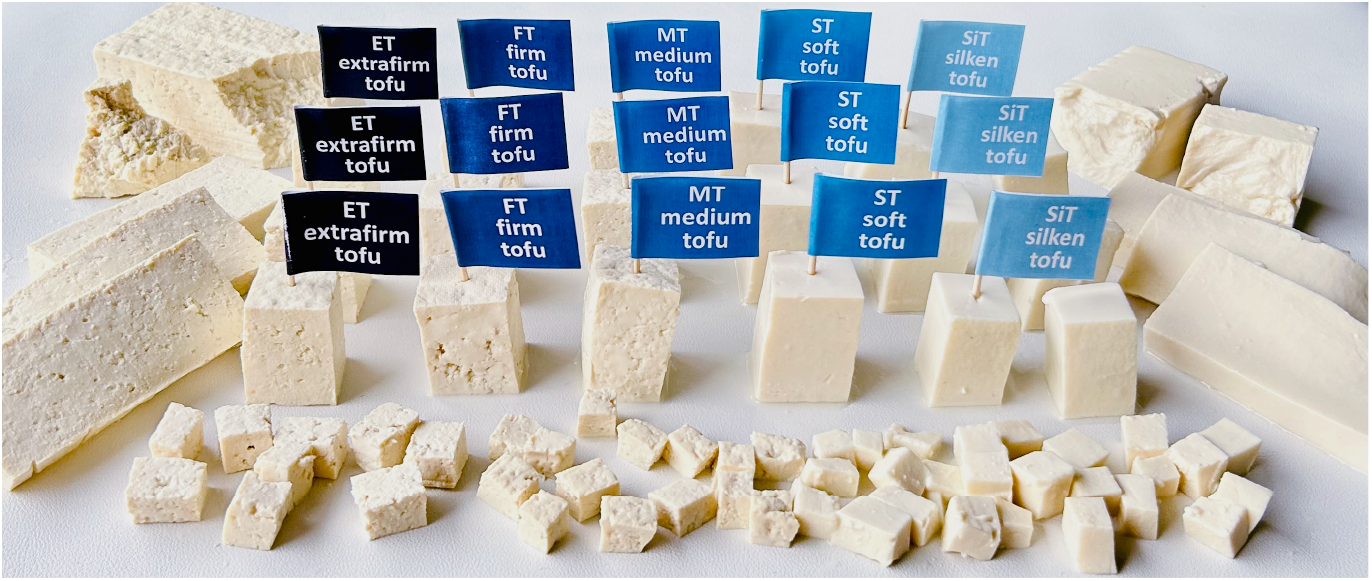
Five different types of tofu. Extra-firm, firm, medium, soft, and silken tofu illustrate how small changes in composition and processing create large differences in mechanical behavior. Tofu combines soy beans with water to form a biphasic protein gel whose stiffness, porosity, and fracture characteristics arise directly from its water content and its coagulation pathway. Whether we make out own tofu at home or obtain it through commercial retail distribution, we encounter the same continuum of textures, from a dense cohesive protein matrix to a nearly fluid-like gel.

For more than half a century, texture profile analysis has defined the gold standard in food science to quantify the textural and sensory attributes of food [26]. In texture profile analysis, mechanical double-compression tests simulate the behavior of chewing [5], we can easily extract its parameters such as stiffness, hardness, cohesiveness, springiness, resilience, and chewiness from the resulting force–time curves [22]. Although texture profile analysis continues to be widely used [60], the lack of standardized testing protocols limits comparability across the different studies and labs [67]. Moreover, the interpretation of texture profile analysis curves remains largely rooted in food science rather than in materials science, even though these tests inherently capture valuable mechanical information.

Mechanical testing allows us to interpret texture through the lens of mechanics and to extract fundamental material properties directly from experimental data [71]. We can quantify material parameters such as stiffness, viscosity, and relaxation time–commonly used in mechanics–from tension, compression, and shear tests to reveal the underlying macrostructural behavior and correlate it to the microstructural architecture [12]. Grounding texture evaluation in the principles of material modeling [77] enables a more rigorous and interpretable mechanistic analysis [72]. This raises the question: Can we characterize the behavior of food through robust constitutive models that reliably reproduce experimental data and confidently predict the effects of different loading conditions?

Food materials form an emerging class of complex, heterogeneous systems that challenge conventional modeling techniques [17]. Tofu, as a hydrated soy protein gel, contains water in three distinct states–tightly bound, immobilized, and freely mobile–each interacting differently with the protein network [45]. While bound and immobilized water contribute to network stability and elastic stiffness, freely mobile water governs fluid redistribution, rate dependence, and syneresis and gives tofu a fundamentally biphasic, poroelastic character [52]. This nonlinear, often rate-dependent response calls for new approaches that are capable of identifying the governing equations directly from experimental data [43]. Recent advances in data-driven mechanics have introduced frameworks that automatically discover constitutive relationships [47] without pre-selecting the specific functional form of the model [28]. Instead of repeatedly fitting and refining candidate models, automated model discovery infers the governing equations in a self-consistent and physics-informed manner [24]. This motivates us to ask: Can we adopt the concept of automated model discovery to discover the models and parameters that best characterize tofu as a minimal benchmark system for soft, structurally engineered foods?

A wide range of data-driven frameworks has emerged in recent years, including unsupervised learning [24] as well as mechanics-informed [3], physics-constrained [38], physics-augmented [39, 40], thermodynamics-based [54], input convex neural networks [2], and input monotonic neural networks [32]. Among these, the framework of constitutive artificial neural networks has proven particularly effective because it hardwires continuum mechanical principles directly into the network architecture [46]. This design ensures that the resulting models remain both physically sound and interpretable [47]. The sparse network structure balances accuracy and interpretability and allows the simultaneous discovery of concise yet descriptive constitutive models and their model parameters.

Automated model discovery has successfully identified constitutive models across diverse applications, including soft biological tissues such as skin [49], arteries [77], the heart [53], and the brain [34, 48], and, most recently, plant-based and animal meats [70]. Most studies have focused on discovering models for the elastic behavior; yet, an accurate representation of realistic material responses–especially at larger loading magnitudes and rates–often requires the inclusion of inelastic effects. Recent extensions of the framework have incorporated viscoelasticity through Prony-series representations [1, 82]. Building on this idea, inelastic constitutive artificial neural networks, extend the formulation by discovering both, the Helmholtz free energy for the elastic behavior and a dual potential for the inelastic behavior [28]. This approach preserves thermodynamic consistency and interpretability, while expanding the range of representable material responses. The general framework of inelastic constitutive neural networks has since been applied to viscoelasticity [29], growth modeling [30], and plasticity [4, 76]. Importantly, inelastic constitutive neural networks autonomously discover whether inelastic effects are necessary, and reduce unnecessary model complexity when the data do not justify it [31].

Motivated by the growing interest in meat alternatives, this study investigates the mechanics and physics of tofu by replacing the traditional texture profile analysis with classical mechanical testing. We perform compression experiments on multiple tofu types with varying water content and analyze the resulting force–time curves to explore the correlation between structure and function. First, we adopt an inelastic constitutive neural network to discover individual constitutive models for each tofu type and identify the dominant mechanisms that govern the material behavior. Then, we postulate that the water content is the single distinguishing feature across all individual models. We thus design a second neural network that takes the water content as input and outputs a feature vector that modulates the inelastic constitutive neural network. This architecture enables a flexible and unbiased mapping between composition and mechanical behavior. It results in a single unified model that provides predictive insight into how changes in formulation–such as water or protein content–affect the texture of tofu, or more generally, of any alternative protein product.

Section 2 describes the mechanical testing. Sections 3 and 4 introduce the physics-based model and the neural network model. Section 5 summarizes our results and Section 6 discusses our main findings and outlines future directions.

## 2. Mechanical testing

We test a total of *n* = 108 samples with three different water contents across three different loading magnitudes at five different rates. Specifically, we test three types of tofu, silken, firm, and extrafirm (Azumaya, Nasoya Foods, Ayer, Massachusetts). Table 1 summarizes their serving sizes, calories, and macronutrient compositions per serving according to their individual nutrition fact labels. We estimate the water mass as the difference between the serving size and the listed masses of fat, carbs, protein, and ash. We assume that tofu fat mainly presents as lipid droplets that do not move freely under mechanical loading, and conclude that the fluid phase of tofu is made up exclusively of the freely moving water. To calculate the mass fraction of the fluid *w*_f_, we divide the water mass, *m*_f_ = {76g, 71g, 70g}, by the total mass per serving of 85g, resulting in *w*_f_ = {0.89, 0.84, 0.82} for the three tofu types. To calculate the volume fraction of the fluid *ν*_f_ = *V*_f_ */*[ *V*_f_ + *V*_s_ ], we divide the volume of the fluid, *V*_f_ = *w*_f_ */ρ*_f_, by the total volume of fluid and solid, *V*_f_ + *V*_s_, with *V*_s_ = [ 1 *− w*_f_ ]*/ρ*_s_, assuming fluid and solid densities of *ρ*_f_ = 1.0 g/ml and *ρ*_s_ = 1.3 g/ml, resulting in *ν*_f_ = {0.92, 0.87, 0.86} for the three tofu types. For each tofu type, we perform four sets of tests, three triple-compression tests at different rates, and one double-compression test. For each set, we test at least *n* = 8 samples.

**Table 1:**
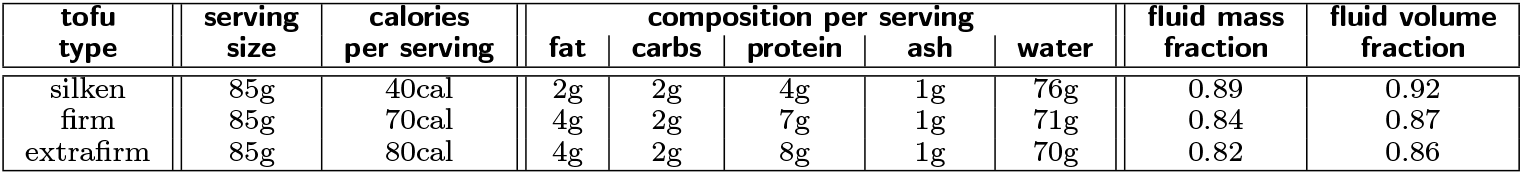
Nutrition facts of silken, firm, and extrafirm tofu. Serving size, calories per serving, macronutrient composition per serving, and fluid mass and volume fractions for the three different tofu types.

| tofu type | serving size | calories per serving | fat | carbs | protein | ash | water | fluid mass fraction | fluid volume fraction |
| --- | --- | --- | --- | --- | --- | --- | --- | --- | --- |
| silken | 85g | 40cal | 2g | 2g | 4g | 1g | 76g | 0.89 | 0.92 |
| firm | 85g | 70cal | 4g | 2g | 7g | 1g | 71g | 0.84 | 0.87 |
| extrafirm | 85g | 80cal | 4g | 2g | 8g | 1g | 70g | 0.82 | 0.86 |

### Sample preparation

To prepare our samples, we first cut the tofu into 10 mm thick blocks and then extract cylindrical specimens of 8 mm diameter using biopsy punches. Figure 2, left, illustrates our sample preparation. To prevent the samples from dehydration, we store them between wetted tissues and test them immediately after preparation.

**Figure 2.**
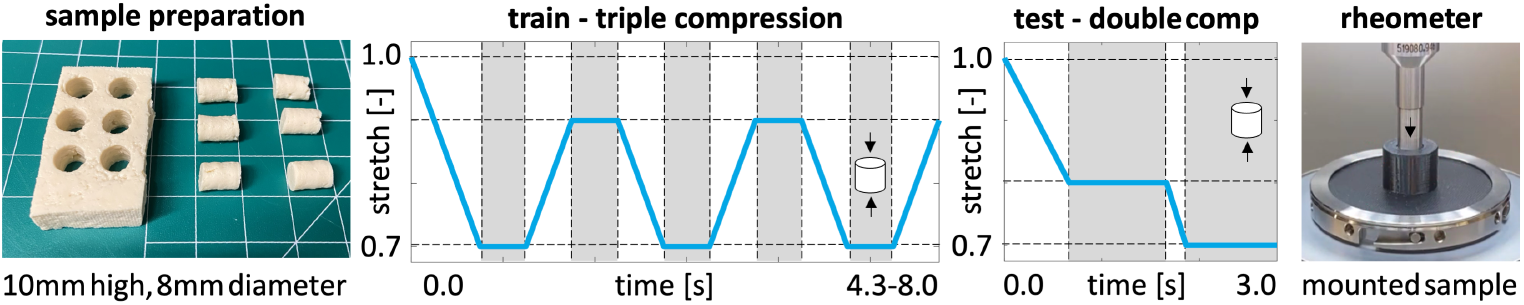
Mechanical testing. Preparation of cylindrical samples; triple compression test for training; double compression test for testing; and rheometer with mounted sample. During triple compression, we load and unload between *λ* = 0.7 and *λ* = 0.9 at slow, medium, and fast stretch rates of 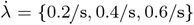 and hold for 0.5s in between. During double compression, we load to *λ* = 0.8 at 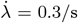, hold for 1.0s, load to *λ* = 0.7 at 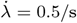, and hold for 1.0s. White regions represent loading and unloading, grey regions represent holding.

### Sample testing

We perform all mechanical tests using an HR20 Discovery Hybrid Rheometer (TA Instruments, New Castle, Delware). Figure 2, right, illustrates a representative sample inside a cylindrical chamber, mounted in the rheometer, to perform compression tests with controlled axial displacements and constrained lateral deformation. Before testing, we calibrate the rheometer, tare the force sensor, and adjust the initial gap between the loading nod and the specimen to ensure contact without exceeding a force of 0.05 N. All experiments follow a displacement-controlled protocol with four distinct loading programs: three triple-compression tests that serve as training data and one double-compression test that serves as test data. During triple compression in Figure 2 middle left, we compress each sample three times to a stretch of *λ* = 0.7, hold for 0.5 s, unload to *λ* = 0.9, and hold for another 0.5 s, at slow, medium, and fast stretch rates of 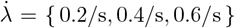. During double compression in Figure 2 middle right, we compress each specimen first to a stretch of *λ* = 0.8 at a rate of 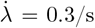, hold for 1.0 s, further compress to *λ* = 0.7 at a rate of 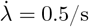, and hold for another 1.0 s.

### Data processing

We record the gap *l* and the force *F* throughout the loading history and smoothen both with a Gaussian filter using the `imgaussfilt` function in Matlab R2022b [55] with a filter width of *σ*_gauss_ = {8, 3, 3} for silken, firm, and extrafirm tofu. We calculate and report the axial stretch *λ* = *l/L* as the ratio between the current gap *l* and the initial gap *L* and the axial second Piola Kirchhoff stress *S* = *F/*(*Aλ*) as the recorded force *F* divided by the initial cross section area *A* = *π* · (4 mm)^2^ and the stretch *λ*.

## 3. Physics-based model

Discovering a physically sound constitutive model that explains the experimental data requires a thermodynamically consistent framework. In this section, we outline the continuum equations that motivate the design of our neural network architecture.

### Kinematics

To characterize the experimentally observed rate dependence and nonlinear relaxation, we adopt a multiplicative decomposition of the deformation gradient ***F*** into elastic and inelastic parts, ***F*** _e_ and ***F*** _i_,

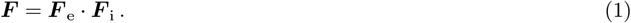

As deformation measures, we introduce the right Cauchy Green deformation tensor ***C*** and its elastic and inelastic counterparts, ***C***_e_ and ***C***_i_,

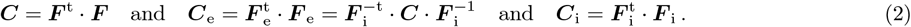

To characterize the isotropic volume-preserving and volume-changing behavior of tofu, we introduce the isochoric invariants 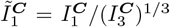 and 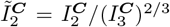 in terms of the classical invariants 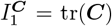 and 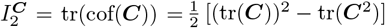, and the volumetric invariant 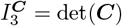 of the right Cauchy Green tensor ***C***,

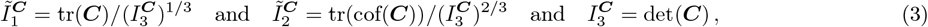

where tr( ○ ) = ( ○ ) : ***I*** denotes the trace, cof( ○ ) = *I*_3_^(○)^ ( ○ )^*−*t^ the cofactor, tr 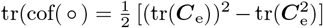 the trace of the cofactor, and det(○ ) the determinant of the second-order tensor (○). The first invariant 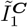 measures the isochoric stretch, the second invariant 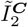 measures the isochoric area change, and the third invariant 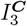 measures the volume change. To characterize the elastic behavior only, we introduce the elastic isochoric invariants 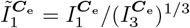 and 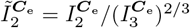 in terms of the classical elastic invariants 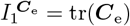 and 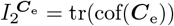, and the volumetric invariant 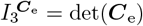, of the elastic right Cauchy Green tensor ***C***_e_,

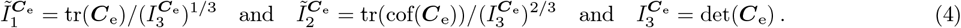

### Elastic potential

We model tofu as an isotropic, inelastic solid and express its total Helmholtz free energy *ψ* as the sum of an equilibrium contribution *ψ*^eq^ for the elastic part that depends on the right Cauchy Green tensor ***C***, or, alternatively, on its invariants 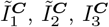, and a non-equilibrium contribution *ψ*^neq^ for the inelastic part that depends on the elastic right Cauchy Green tensor ***C***_e_, or on its invariants 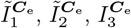,

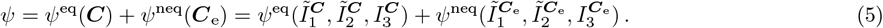

To ensure thermodynamic consistency, we evaluate the Clausius–Planck inequality, 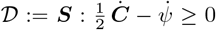, where ***S*** is the second Piola Kirchhoff stress and the dot denotes the material time derivative,

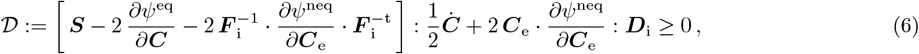

where 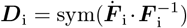 is the inelastic rate of deformation tensor. From the Clausius-Planck inequality (6), we obtain the definitions of the second Piola Kirchhoff stress ***S*** := 2 *∂ψ/ ∂****C***,

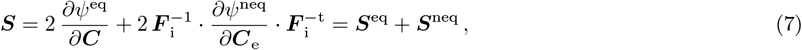

its equilibrium part ***S***^eq^ := 2 *∂ψ*^eq^*/ ∂****C***,

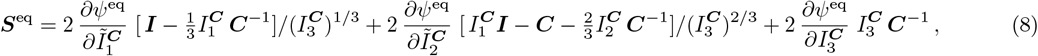

and its non-equilibrium part 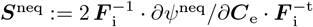,

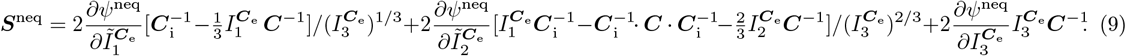

In Section 4, we reverse-engineer two constitutive neural networks, one for the equilibrium potential *ψ*^eq^ and one for the non-equilibrium potential *ψ*^neq^, from which we derive the equilibrium and non-equilibrium stresses, ***S***^eq^ and ***S***^neq^, according to equations (8) and (9).

### Dissipative potential

From the Clausius-Planck inequality (6), we motivate the Mandel stress **Σ** as the driving force for dissipative processes, with **Σ** := 2 ***C***_e_ · *∂ψ*^neq^*/ ∂****C***_e_,

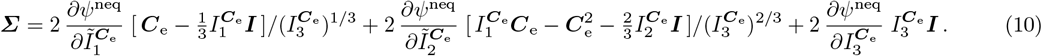

To characterize the dissipative volume-changing and volume-preserving behavior of tofu, we introduce the volumetric invariant 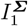 and the deviatoric invariants 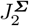 and 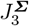 of the Mandel stress **Σ**,

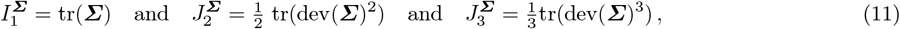

with derivatives 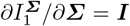 and 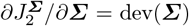 and 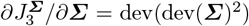, where tr( ○ ) = ( ○ ) : ***I*** denotes the trace and dev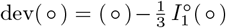 denotes the deviator of a second-order tensor ( ○ ). The first invariant 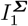 measures the hydrostatic stress, the second invariant 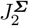 measures the magnitude of shear, and the third invariant 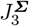 measures the magnitude and mode of shear. To characterize the evolution of the inelastic deformation, we introduce the dissipative potential *ϕ* that depends on the Mandel stress **Σ** or, alternatively, on its invariants 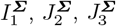,

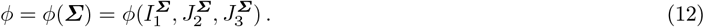

To a priori satisfy the reduced Clausius–Planck inequality,

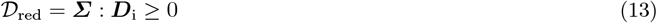

we postulate that the dissipative potential *ϕ* is convex, non-negative, and zero at the origin and define the inelastic rate of deformation tensor as ***D***_i_ := *∂ϕ/ ∂***Σ**,

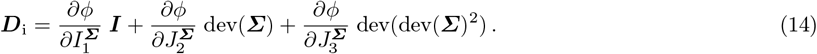

The inelastic rate of deformation tensor ***D***_i_ introduces the evolution of the inelastic right Cauchy Green tensor, 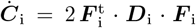, which we now discretize in time to calculate the inelastic right Cauchy Green tensor ***C***_i_ and its elastic counterpart ***C***_e_.

### Time discretization

To advance the governing equations in time, from the previous time point *t*^n^ with known deformation gradient ***F*** ^n^, inelastic tensor 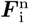, right Cauchy Green tensor 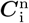, and rate of deformation tensor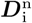, to the new time point *t*^n+1^ = *t*^n^ + Δ*t* with time increment Δ*t*, we adopt an explicit time integration scheme using the exponential update of the inelastic right Cauchy Green tensor,

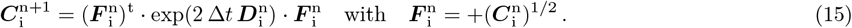

In Section 4, we design a neural network for the dissipative potential *ϕ* from which we derive the inelastic rate of deformation tensor ***D***_i_, the inelastic right Cauchy Green tensor ***C***_i_, and the inelastic part of the deformation gradient ***F*** _i_ at the new time point *t*^*n*+1^.

### Features

In many engineering applications, additional state variables or fields such as the density, temperature, or water content may affect the material behavior. To account for these effects, we extend the constitutive framework by introducing an input feature vector ***α*** that can modify the elastic and dissipative potentials,

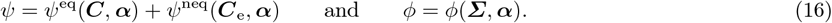

Here, we acknowledge the biphasic nature of tofu and propose a single scalar-valued input feature, the water content *α* = *n*^F^. In Section 4, we design a feature network that we couple to the networks for the equilibrium potential *ψ*^eq^, the non-equilibrium potential *ψ*^neq^, and the dissipative potential *ϕ* to explore the dependencies of tofu on the water content.

### Kinematics of finite inelastic confined cylindrical compression

Our experiment considers a cylindrical sample under confined compression in the *z*-direction. Its principal stretches in cylindrical coordinates, *λ*_*r*_ = 1, *λ*_*θ*_ = 1, *λ*_*z*_ = *λ*, define the total deformation gradient ***F*** that we multiplicatively decompose into elastic and inelastic parts, ***F*** _e_ and ***F*** _i_,

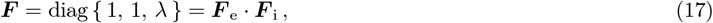

with the Jacobian, *J* = det(***F*** ) = *λ*_*r*_ *λ*_*θ*_ *λ*_*z*_ = *λ*, defining the volumetric change of the sample. We assume that the inelastic deformation is isotropic and volume preserving, *J*_i_ = det(***F*** _i_) = 1, and that all volumetric changes are associated with the elastic deformation only, *J*_e_ = det(***F*** _e_) = *λ* [33, 65, 69] *such* that elastic and inelastic parts of the deformation gradient are

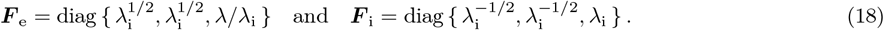

These explicit definitions introduce the right Cauchy Green tensor ***C*** and its elastic counterpart ***C***_e_,

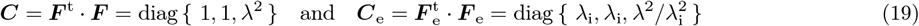

with invariants

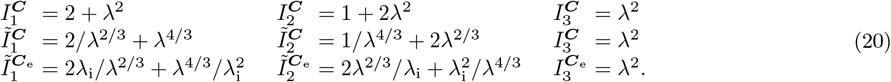

We note that, consistent with axisymmetric confined compression, all tensors are diagonal in the principal directions {*r, θ, z }*, i.e., there are no shear components, and the radial and circumferential components are identical since the deformation is axisymmetric. For this special case of inelastic confined cylindrical compression, we can explicitly calculate the equilibrium stress in the axial direction,

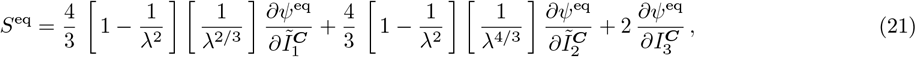

and the non-equilibrium stress in axial direction,

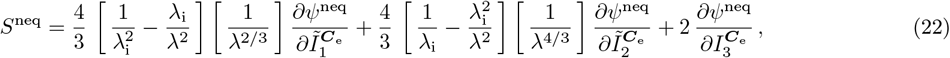

solely in terms of total and inelastic stretches, *λ* and *λ*_i_, and the derivatives of the elastic potentials, *ψ*^eq^ and *ψ*^neq^, that we will now discover using constitutive neural networks.

## 4. Neural network model

To discover the model and parameters that best describe our experimental data, we introduce a combination of four constitutive neural networks for the equilibrium potential *ψ*^eq^ [47], the non-equilibrium potential *ψ*^neq^ [30], the dissipative potential *ϕ* [31], and the weight vector *F* ^max^.This architecture combines two feed forward neural networks for the equilibrium and non-equilibrium potentials with a recurrent neural network for the dissipative potential within the framework of inelastic constitutive neural networks, and, if desired, combines the resulting overall architecture with a parallel feed forward network for the weight vector. It takes the known deformation state ***F*** ^n^ and the internal variables 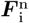 from the previous time point *t*^n^ as input, integrates them in time using the exponential update (15), stores the internal variables 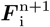, and outputs the second Piola Kirchhoff stress ***S***^n+1^ at the new time point *t*^n+1^.

### Elastic networks

To model the equilibrium potential *ψ*^eq^ [47] and the non-equilibrium potential *ψ*^neq^ [31], we introduce two structurally similar but input-independent feed forward networks. The equilibrium network uses the invariants of the total right Cauchy Green tensor 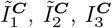 from equation (3) [48, 53, 61] and the non-equilibrium network uses the invariants of the elastic right Cauchy Green tensor 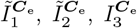 from equation (4) [28, 29, 30] as inputs. For both networks, we ensure expressiveness and interpretability by adopting a sparse network architecture with two hidden layers, where the activation functions *f* ^*I*^ of the first layer are the first and second powers and the activation functions *f* ^*II*^ of the second layer are the identity, the exponential function, and the natural logarithm,

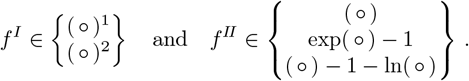

Figure 3 visualizes the architecture of both elastic networks that a priori satisfy the thermodynamic requirements of polyconvexity, growth, and normalization [4, 31, 47]. The equilibrium and non-equilibrium networks each contain 14 trainable parameters, which are visualized in Figure 3 through solid lines. We constrain all weights to be non-negative and add Lasso regularization to the weights of the final layer. Importantly, although the two feed forward networks for the equilibrium and non-equilibrium potentials are input-independent, they are coupled through their derivatives.

**Figure 3.**
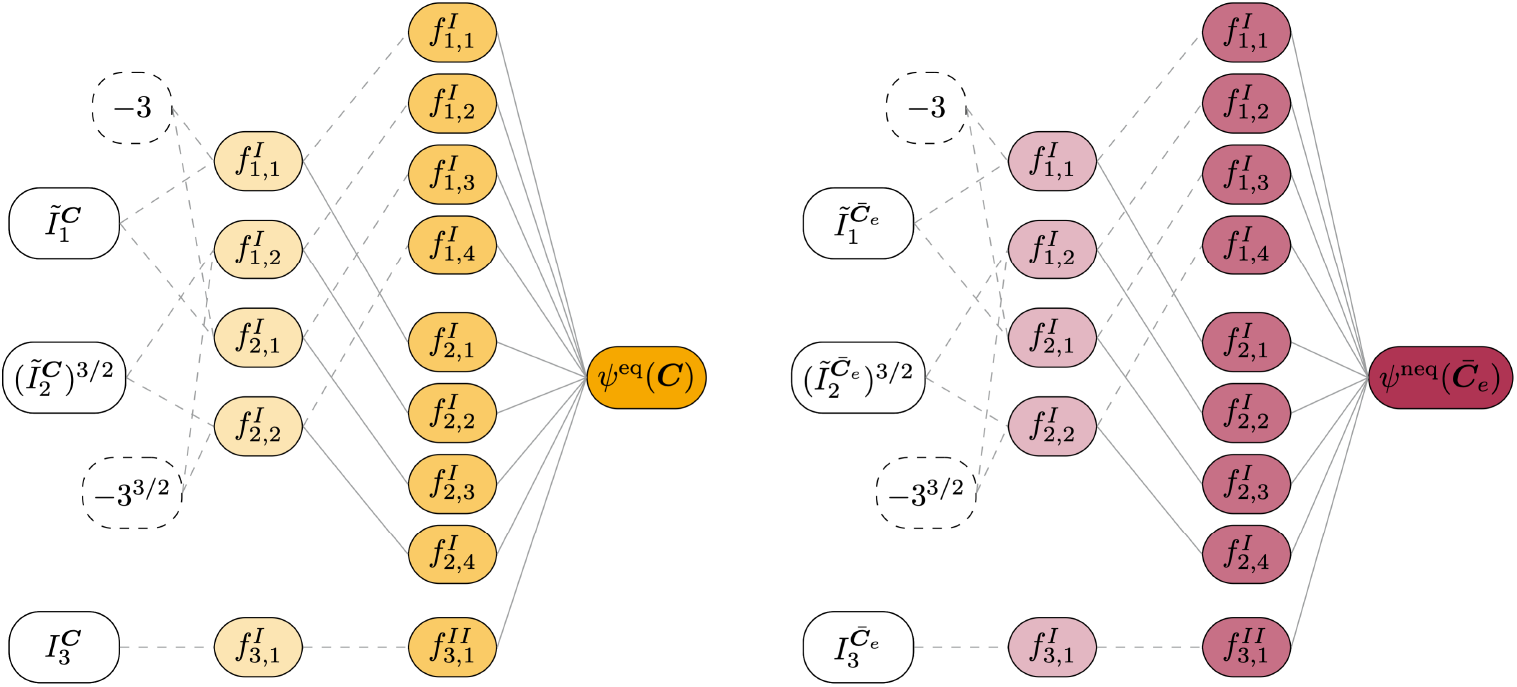
Elastic networks. The elastic networks have two hidden layers with 5 and 9 nodes per layer. The feed forward networks take the invariants of the total right Cauchy Green tensor 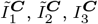, left, or of the elastic right Cauchy Green tensor 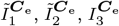, right, as input; the first layer applies first or second powers, the second layer applies the identity, exponential function, or natural logarithm; the networks learn the equilibrium or non-equilibrium potentials *ψ*^eq^, left, or *ψ*^neq^, right, as output. Activation functions 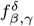 are labeled by activation type *β*, node index *γ*, and layer index *δ*; solid lines denote trainable weights, dashed lines indicate fixed weights equal to one, and dashed ellipses denote constant, non-trainable biases.

### Dissipative network

To model the dissipative potential *ϕ* [31], we introduce a recurrent neural network that uses the invariants of the Mandel stress 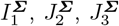 from equation (11) as input. Specifically, to maintain consistent physical units, we take their first, second, and third roots as input. We adopt a fully connected network architecture with three hidden layers, where the activation functions *f* ^*I*^ of the first layer are trainable powers *p* of the absolute input values |( ○ )|, and the activation functions *f* ^*II*^ and *f* ^*III*^ of the second and third layers are the maximum or exponential functions,

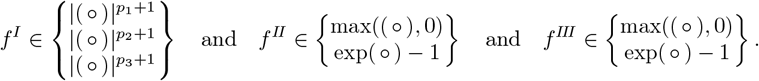

Figure 4 visualizes the architecture of the dissipative network that a priori satisfies thermodynamic consistency by remaining non-negative, convex, and zero-valued at equilibrium to ensure that the reduced dissipation inequality 𝒟 _red_ *≥* 0 (13) is satisfied. In the first layer, we smoothen the absolute-value functions | (○) |near the origin for differentiability, constrain the exponents *p*_1_, *p*_2_, *p*_3_ to be non-negative, and allow the network weights to take positive or negative values [31]. In the second and third layers, we apply additional non-positive biases to the maximum functions and constrain the weights to be non-negative. To guarantee non-negativity, we introduce a final layer that mirrors the third layer, but uses the maximum function without additional weights. After evaluating multiple network configurations with varying numbers of neurons per layer, we select the architecture that achieves the best performance in a preliminary study: a three-layered network with {18, 8, 8} nodes per layer. The resulting dissipative network has a total number of 281 trainable parameters. Appendix B provides additional sensitivity analyses for varying network architectures and varying activation functions.

**Figure 4.**
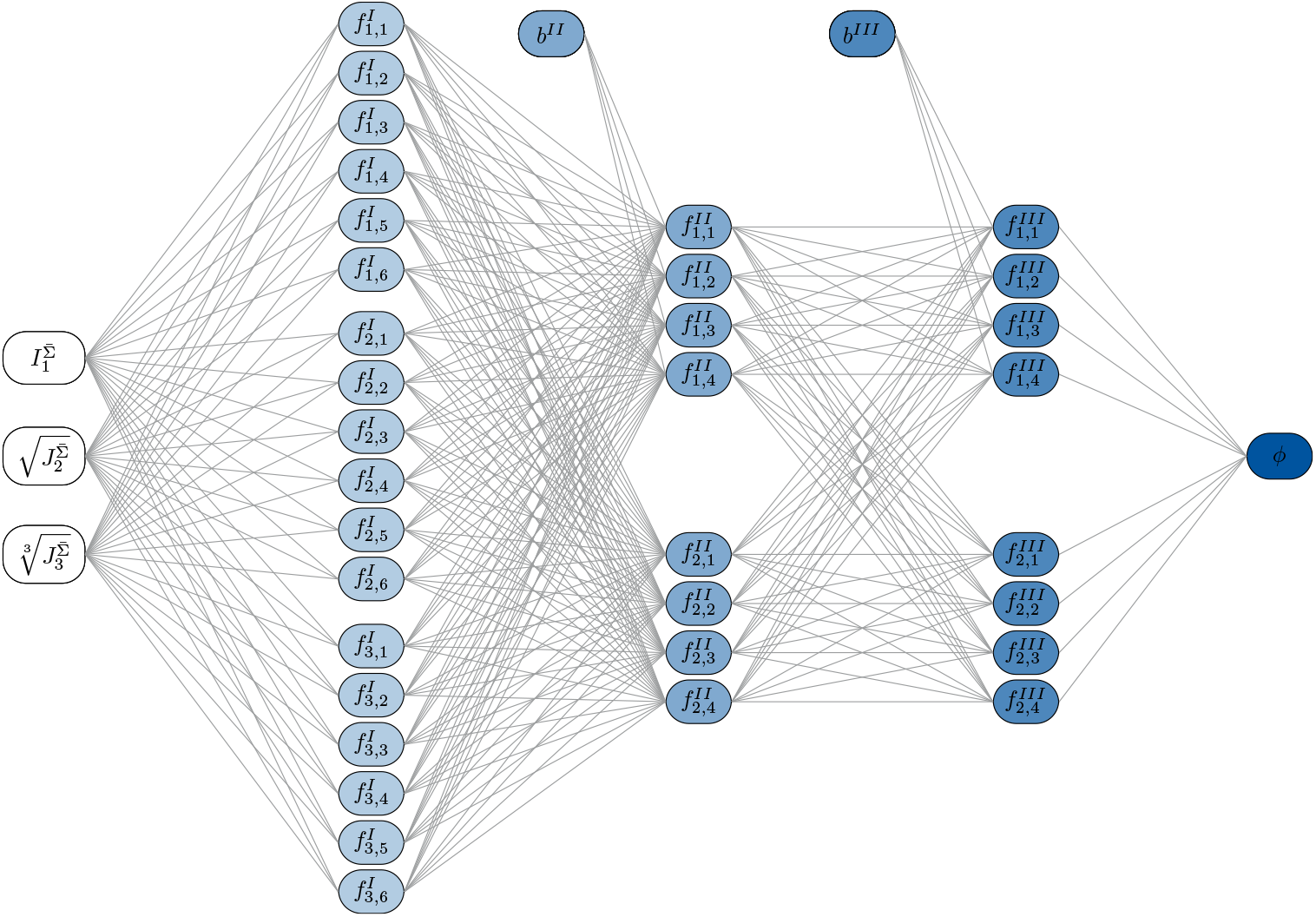
Dissipative network. The dissipative network has three hidden layers with 18, 8, and 8 nodes per layer. It take the invariants of the Mandel stress 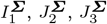 as input; the first layer applies trainable powers *p* to the absolute input values, the second and third layers apply the maximum or exponential functions; the network learns the dissipative potential *ϕ* as output. Activation functions 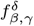 are labeled by activation type *β*, node index *γ*, and layer index *δ*; biases *b*^*δ*^ are applied to the maximum functions in the second and third hidden layers.

### Feature network

To model the impact of the feature vector on the mechanical response, we introduce a feed forward network that uses the water content as a single scalar-valued input feature *α*. We adopt a fully connected network architecture with three hidden layers, where the activation functions *F* ^*I*^, *F* ^*II*^, *F* ^*III*^ are the maximum, identity, hyperbolic tangent, and softplus functions and the output of the final layer *F* ^max^ applies the maximum function,

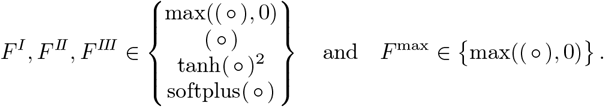

Figure 5 schematically visualizes the architecture of the feature network. To a priori ensures non-negative, non-zero-valued outputs *F* ^max^, we add biases to all maximum functions. After evaluating different network architectures, we select feature networks with {14,15,11} nodes per activation function per layer for the equilibrium potential, {13,17,15,9} for the non-equilibrium potential, and {20,6,6} for the dissipative potential. The resulting feature network adds a total number of 19,494 trainable parameters. The number of nodes in the final layer *n*^max^ matches those of the mechanical network to which it is coupled, with *n*^max^ = 9 nodes for the elastic networks in Figure 3 and *n*^max^ = 8 nodes for the dissipative network in Figure 4.

**Figure 5.**
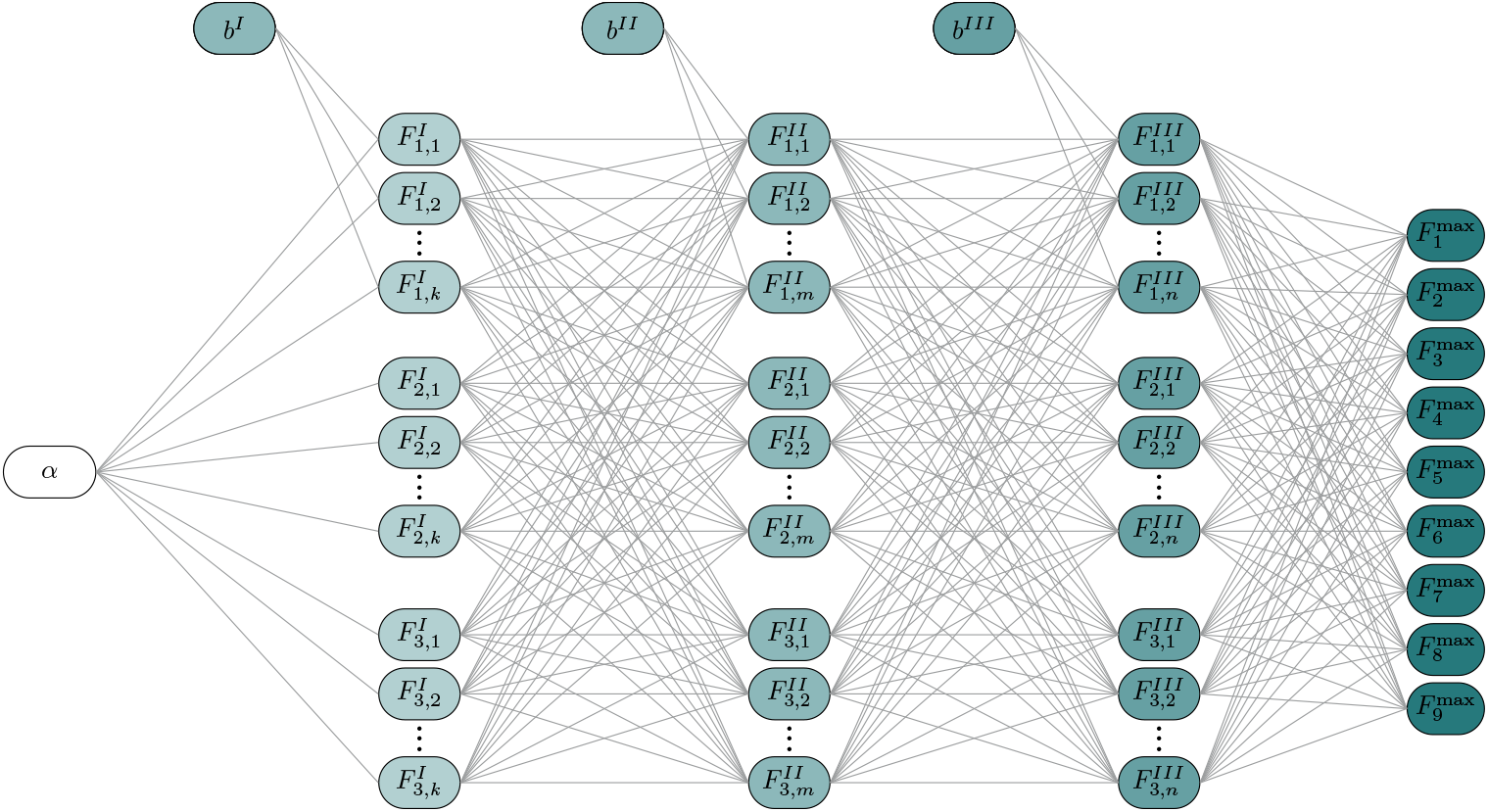
Feature network. The feature network has three hidden layers with *k ×* 4 nodes per layer. It takes the feature *α*, in our case the scalar-valued water content, as input; the three hidden layer apply the maximum, identity, hyperbolic tangent, and softplus functions; the output of the final layer applies the maximum function; the network learns the feature weight vector *F* ^max^ as output. Activation functions 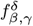 are labeled by activation type *β*, node index *γ*, and layer index *δ*; biases *b*^*δ*^ are applied to all maximum functions.

Figure 6 illustrates the coupling between one of the three mechanical networks and the feature network. The mechanical networks from Figure 3 or 4 receive a strain- or stress-like input ***A*** and the feature network from Figure 5 receives a feature-like input ***α***. The networks are multiplicatively coupled by summing up the product of the nodes in their final layers to obtain feature-weighted potential *Ψ*,

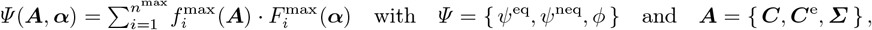

where the potential *Ψ* is either the equilibrium or non-equilibrium potential *ψ*^eq^ or *ψ*^neq^, or the dissipative potential *ϕ*, and the input ***A*** is either the total or the elastic right Cauchy Green tensor ***C*** or ***C***_e_ or the Mandel stress **Σ**. This multiplicative coupling ensures that the combined output is non-negative and remains zero for zero-valued mechanical outputs. Multiplicative feature weighting inherently preserves polyconvexity and other thermodynamic requirements.

**Figure 6.**
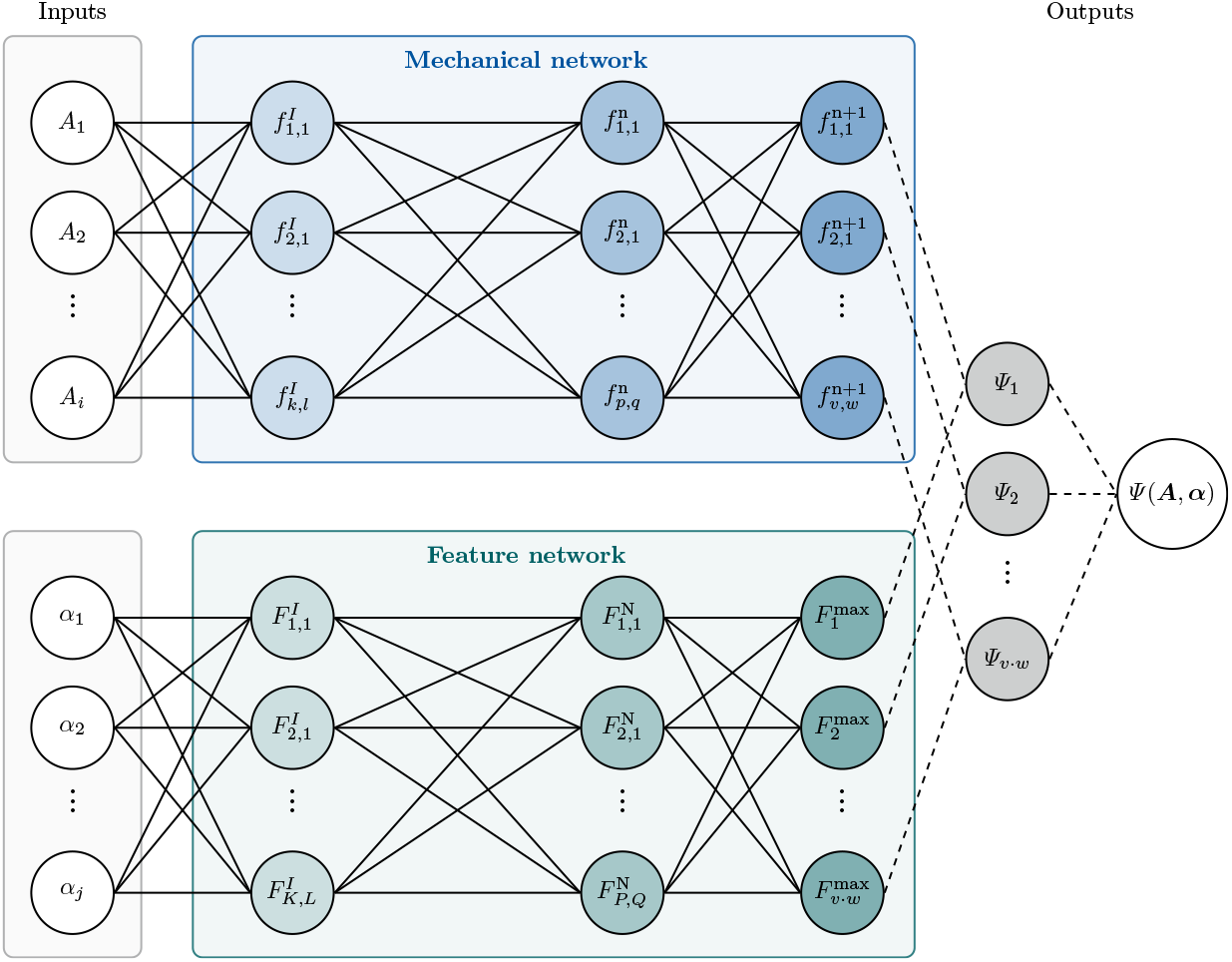
Coupling of mechanical and feature networks. The mechanical network from Figure 3 or 4 receives a strain- or stress-like input ***A*** and the feature network from Figure 5 receives a feature-like input ***α***. The networks are multiplicatively coupled in the nodes of their final layers. The sum of the final layer defines the network output, the feature-weighted mechanical potential *Ψ* (***A, α***). Activation functions 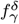 and 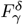 of the mechanical and feature networks are labeled by node index *γ* and layer index *δ*.

### Network training

To discover the best model and parameters to characterize tofu, we train and test our networks using the experimental data from Section 2. We introduce the loss function as the mean squared error between the model stress *S* and the experimental stress *Ŝ* in the direction of loading, averaged across all *n*_exp_ experiments and all *n*_data_ data points. To induce sparsity and improve generalization, we supplement the loss with *L*_1_-regularization or Lasso and *L*_2_-regularization or ridge regression across all *n*_*w*_ weights, where *α*_1_ and *α*_2_ are the regularization coefficients. The total loss function reads

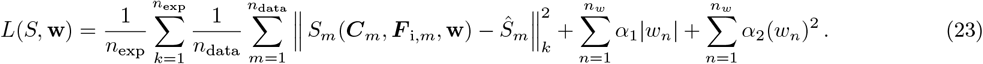

We perform two types of model discovery: First, we analyze *each tofu type independently* by using only the elastic and dissipative networks to discover *three individual tofu models* for each different tofu type. Then, we analyze *all three tofu types combined* by integrating the elastic and dissipative networks with the feature network to discover a *single unified tofu model* with the water content as input feature. Appendix B provides additional sensitivity analyses for varying regularization coefficients.

### Discovering three individual tofu models

First, we train the elastic and dissipative networks to discover individual tofu models using the stress–strain data from Section 2. We neglect water-content dependency and discover three independent material models by simultaneously learning the equilibrium and non-equilibrium potentials *ψ*^eq^ and *ψ*^neq^ and the dissipative potential *ϕ*. We use the triple-compression tests at three different strain rates as training data and the double-compression tests at intermediate strain rates as test data. To capture history-dependent effects, we adopt a staggered training strategy that progressively increases the batch size and gradually introduces additional training data along the load path. We adopt the Adam optimizer with a learning rate of 10^*−*3^ and a clip norm of 10^*−*3^. The training proceeds in two phases: In the first phase, we only include the first *n* = 50 data points of the first cycle of the triple-compression test. In the second phase, we include all cycles and apply a regularization with *α*_1_ = 0.0001 and to all layers of all networks and *α*_2_ = 0.0001 to the first layer of the dissipative network. The weights and biases from the first phase initialize the second phase to ensure continuity and convergence throughout the process.

### Discovering a single unified tofu model

Then, we train the feature networks to discover a unified tofu model for the stress–strain data from Section 2. We incorporate the water-content as a scalar-valued input feature and discover a single material model by learning feature-weighted terms of the equilibrium, non-equilibrium, and dissipative potentials, *ψ*^eq^ and *ψ*^neq^ and *ϕ*. Yet, unlike the elastic and dissipative networks, we do not train the feature network directly on the experimental data. Instead, we train on the *outputs of the discovered models*. We generate synthetic training data by evaluating our three discovered models for various sets of invariants 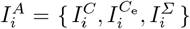 of our network input ***A*** = { ***C, C***_e_, **Σ** }, the total and elastic right Cauchy Green tensors ***C*** and ***C***_e_ and the Mandel stress **Σ**. According to equations (8), (9), and (14), the derivatives of the potentials *Ψ* = *{ ψ*^eq^, *ψ*^neq^, *ϕ }* with respect to these invariants,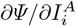, determine both the stress response and the inelastic flow direction { ***S***^eq^, ***S***^neq^, ***D***_i_ }. We train the combined feature–mechanical network on these derivatives, which contain the essential physical information.

Figure 7 illustrates how we generate our synthetic training data 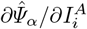: We evaluate each of the three discovered network potentials 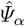 for a series of invariants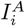, where *α* = *{ n*_silken_, *n*_firm_, *n*_extra_ } is the water content of silken, firm, and extrafirm tofu and *i* = { 1, 2, 3 } are the three isotropic invariants. The derivatives of the potentials with respect to the invariants, 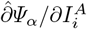, represent the training data that we seek to approximate by the unified model *Ψ* (***A***, *α*, **w**) of the feature-mechanical network that takes both the invariants of ***A*** and the feature *α* as input. During training, the networks learn the weights **w** by minimizing the loss function that measures the error between the derivatives of the unified model 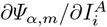 and the derivatives of the synthetic training data 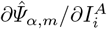 at *m* = 1,…, *n*_data_ data points,

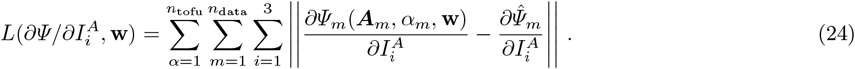

**Figure 7.**
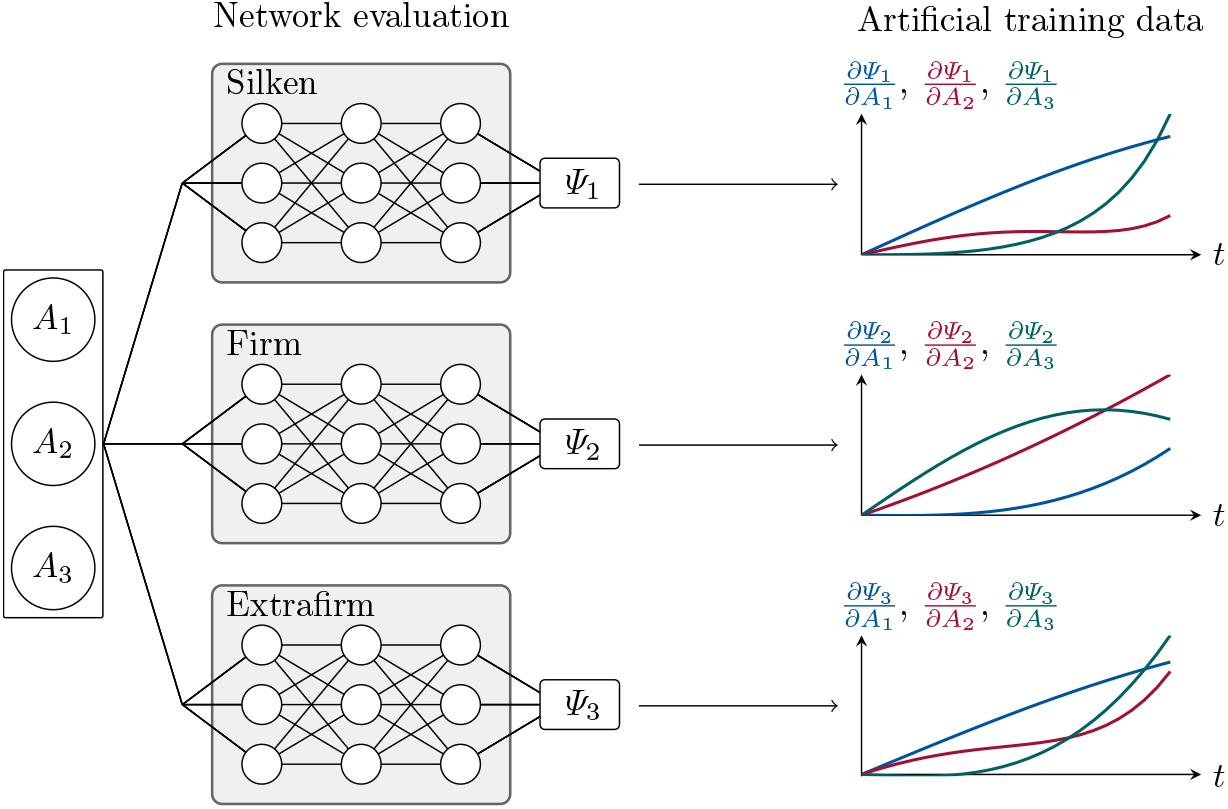
Generating synthetic training data. After discovering the best models and parameters for for silken, firm, and extrafirm tofu by training the equilibrium and non-equilibrium networks in Figures 3 and the dissipative network in Figure 4 on the experimental data, we generate synthetic data to train the coupled mechanical and feature networks in Figure 6. We evaluate the three tofu models for sets of invariants 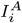, and their derivatives 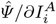 provide the training data for the unified feature–mechanical model.

We adopt the Adam optimizer and train the equilibrium network for 5,000 epochs using 500 training samples with a learning rate of 10^*−*3^ and a clip norm of 10^*−*3^, and the non-equilibrium and dissipative networks for 15,000 and 10,000 epochs with a learning rate of 10^*−*4^. This approach eliminates the need to retrain the entire recurrent network on experimental stress–strain pairs and significantly accelerates the learning process. Importantly, it produces a *single tofu model* that captures tofu types of different water content within a unified constitutive framework.

## 5. Results

### Mechanical testing

Figures 8 and 9 summarize the applied stretch and stress histories *λ*(*t*) and *S*(*t*) during the triple-compression and double-compression tests. Each row corresponds to one tofu type—silken, firm, and extrafirm—and each column represents a different loading condition. The first three columns show the triple-compression tests at different strain rates and the final column shows the double-compression tests. The curves display high reproducibility across all tofu types and loading conditions and confirm that we can robustly record stretch and stress histories at different stretch magnitudes and rates. However, especially at higher strain rates, we observe slight variations in the applied stretch profiles resulting in slight inconsistencies in the stress profiles, which complicate direct averaging of the curves. We thus report each curve as an independent measurement. As a general trend, we observe a non-linear stress response that is sensitive to both strain rate and tofu type: extrafirm tofu is the stiffest, closely followed by firm tofu, and silken tofu is the softest. The stiffness decreases with increasing water content, consistent with the transition from solid-like to fluid-like behavior.

**Figure 8.**
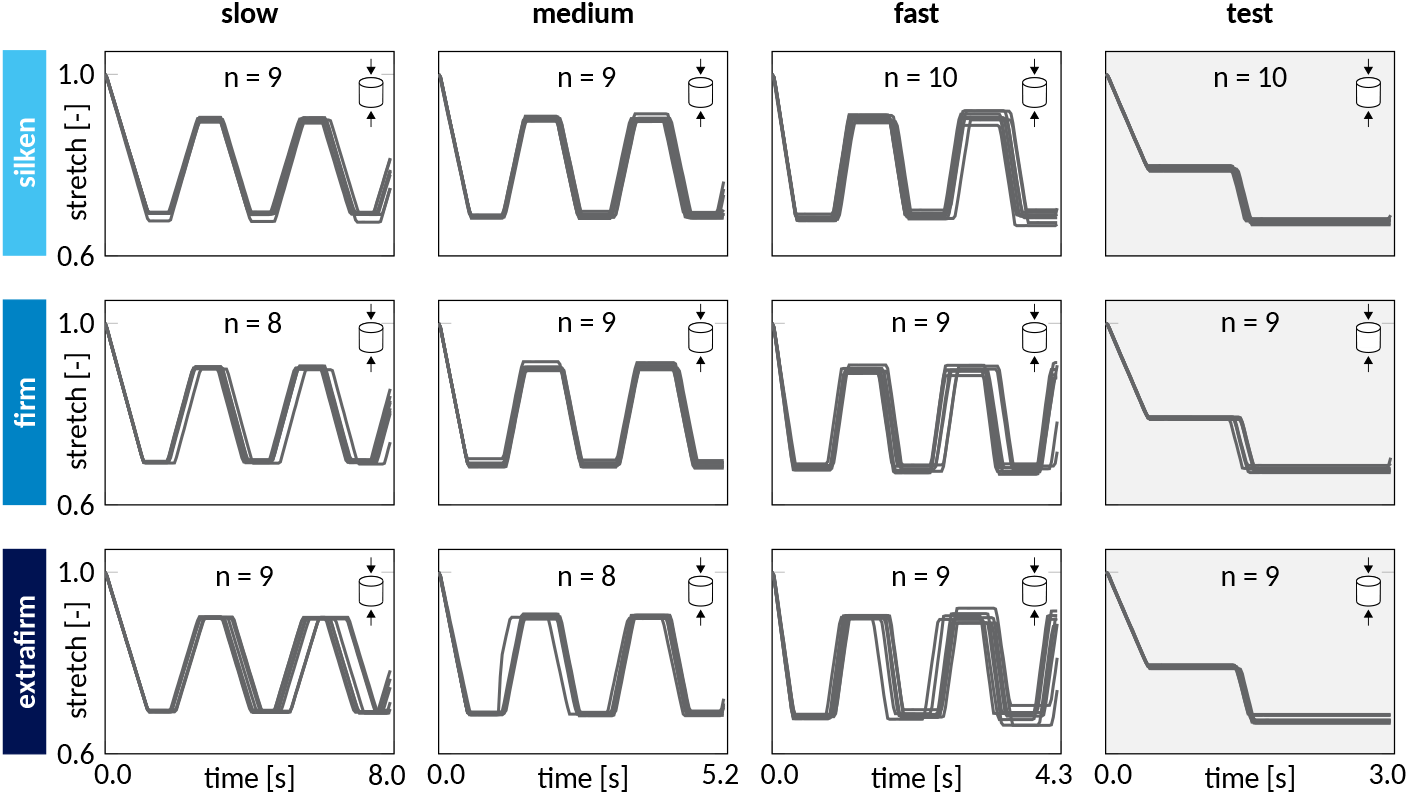
Mechanical testing - Experimentally applied stretches. Stretch histories *λ*(*t*) during triple-compression and double-compression tests for silken (top), firm (middle), and extrafirm (bottom) tofu. The first three columns show the triple-compression tests at slow, medium, and fast strain rates of 20%*/*s, 40%*/*s, and 60%*/*s (from left to right), where each specimen is loaded to a stretch of 0.7 and unloaded to 0.9. The last column shows the double-compression tests at a strain rates of 30%*/*s to a stretch of 0.8 followed by a strain rate of 50%*/*s to 0.7.

**Figure 9.**
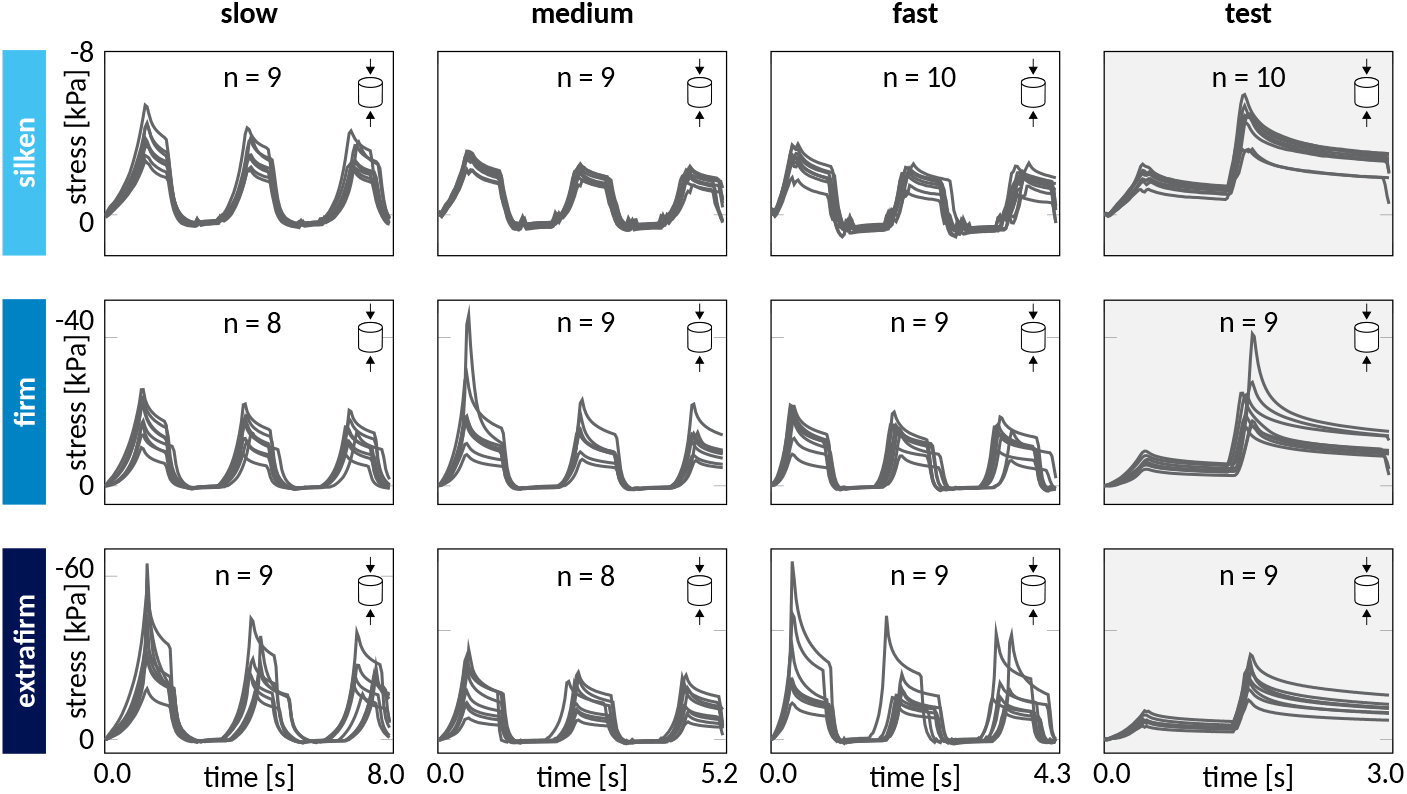
Mechanical testing - Experimentally recorded stresses. Stress histories *S*(*t*) during triple-compression and double-compression tests for silken (top), firm (middle), and extrafirm (bottom) tofu. The first three columns show the triple-compression tests at slow, medium, and fast strain rates of 20%*/*s, 40%*/*s, and 60%*/*s (from left to right), where each specimen is loaded to a stretch of 0.7 and unloaded to 0.9. The last column shows the double-compression tests at a strain rates of 30%*/*s to a stretch of 0.8 followed by a strain rate of 50%*/*s to 0.7.

### Discovering three individual tofu models

Figure 10 compares the computationally predicted stresses in blue and the experimentally recorded stresses in gray. The rows correspond to silken, firm, and extrafirm tofu, and the columns represent distinct loading cases with varying stretch magnitudes and rates. The three discovered models and parameter sets for each tofu type accurately reproduce the experimental trends: For the training data in the first three columns, the modeled stresses generally fall within the measured stress ranges and accurately capture the mean behavior during loading, holding, and unloading, with coefficients of determination ranging from 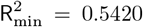to 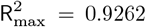. Appendix A provides additional details about the errors. For the test data in the last column, the silken tofu model accurately reproduces the first peak and the relaxation behavior, but slightly underestimates the second peak; the firm tofu model displays a similar trend; and the extrafirm model aligns closely with all experimental curves. The test results show a decrease in the coefficient of determination, but the overall values of 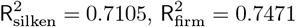 and 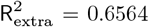 suggest good fits for all three tofu types. Across all tofu types, the discovered models capture a rate dependent behavior: the higher the strain rate, the stiffer the response and the larger the peak stress. In a few cases, the slowest strain rate generates higher peak stresses than the intermediate rate, but overall, the discovered models correctly reproduce strain-rate stiffening.

**Figure 10.**
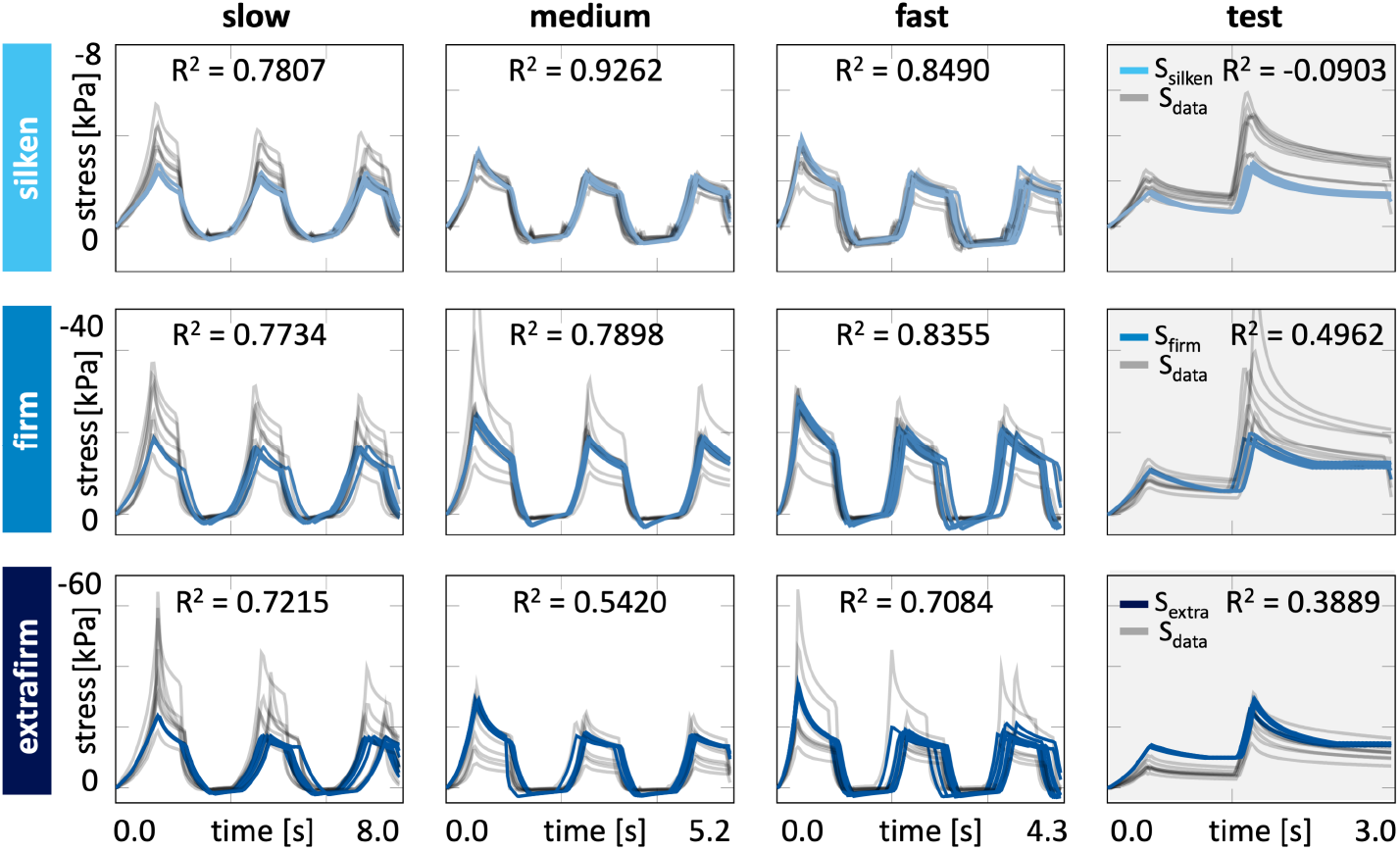
Model discovery - Experimentally recorded and computationally predicted stresses of the three individual tofu models. Stress histories *S*(*t*) for silken (top), firm (middle), and extrafirm (bottom) tofu. The first three columns show the triple-compression tests used as training data. The last column shows double-compression tests used as test data. Experimental data are plotted in gray, model predictions for silken, firm, and extrafirm tofu in light to dark blue; R^2^ values indicate the goodness of fit.

For silken tofu, the discovered model features a quadratic second-invariant term and a logarithmic third-invariant term for the equilibrium potential and a linear second-invariant term for the non-equilibrium potential,

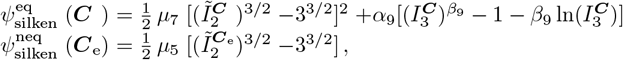

with network weights *µ*_7_ = 0.158 kPa, *α*_9_ = 0.052 kPa, *β*_9_ = 5.507 for the equilibrium part and *µ*_5_ = 1.179 kPa for the non-equilibrium part that have the physical unit of stress and a clear physical interpretation as stiffness-like parameters. For firm tofu, the discovered model features a quadratic first-invariant term and a linear second-invariant term for the equilibrium potential and a linear second-invariant term and a logarithmic third-invariant term for the non-equilibrium potential,

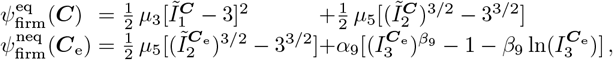

with network weights *µ*_3_ = 5.250 kPa, *µ*_5_ = 0.136 kPa for the equilibrium part and *µ*_5_ = 4.952 kPa, *α*_9_ = 0.867 kPa, *β*_9_ = 1.439 for the non-equilibrium part, again, all with a clear physical interpretation. For extrafirm tofu, the discovered model features a quadratic second-invariant term and a logarithmic third-invariant term for the equilibrium potential and a linear and a quadratic second-invariant term for the non-equilibrium potential,

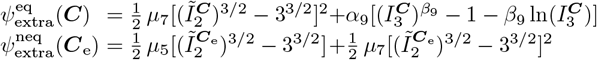

with *µ*_7_ = 0.151 kPa, *α*_9_ = 0.103 kPa, *β*_9_ = 4.920 for the equilibrium part and *µ*_5_ = 8.160 kPa, *µ*_7_ = 2.735 kPa for the non-equilibrium part. Notably, the most prevalent invariant across all six potentials is the second invariant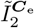 featured in seven terms, followed by the third invariant 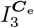 in three terms, and the first invariant 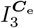 in only one term. None of the models features any of the four exponential terms, neither linear nor quadratic, neither in the first nor in the second invariant, and none of the models features a quadratic first-invariant term.

The dissipative part of our model discovery uses the dissipative network in Figure 4 to discover the dissipative potential *ϕ*. Figures 11–13 display the three discovered potentials for silken, firm, and extrafirm tofu. The colors represent the magnitude of the final weights and biases and enable direct interpretation of active pathways. For silken tofu in Figure 11, the dissipative potential is driven by both the second and third deviatoric invariants 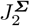 and 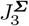,

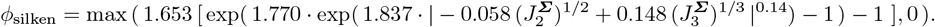

**Figure 11.**
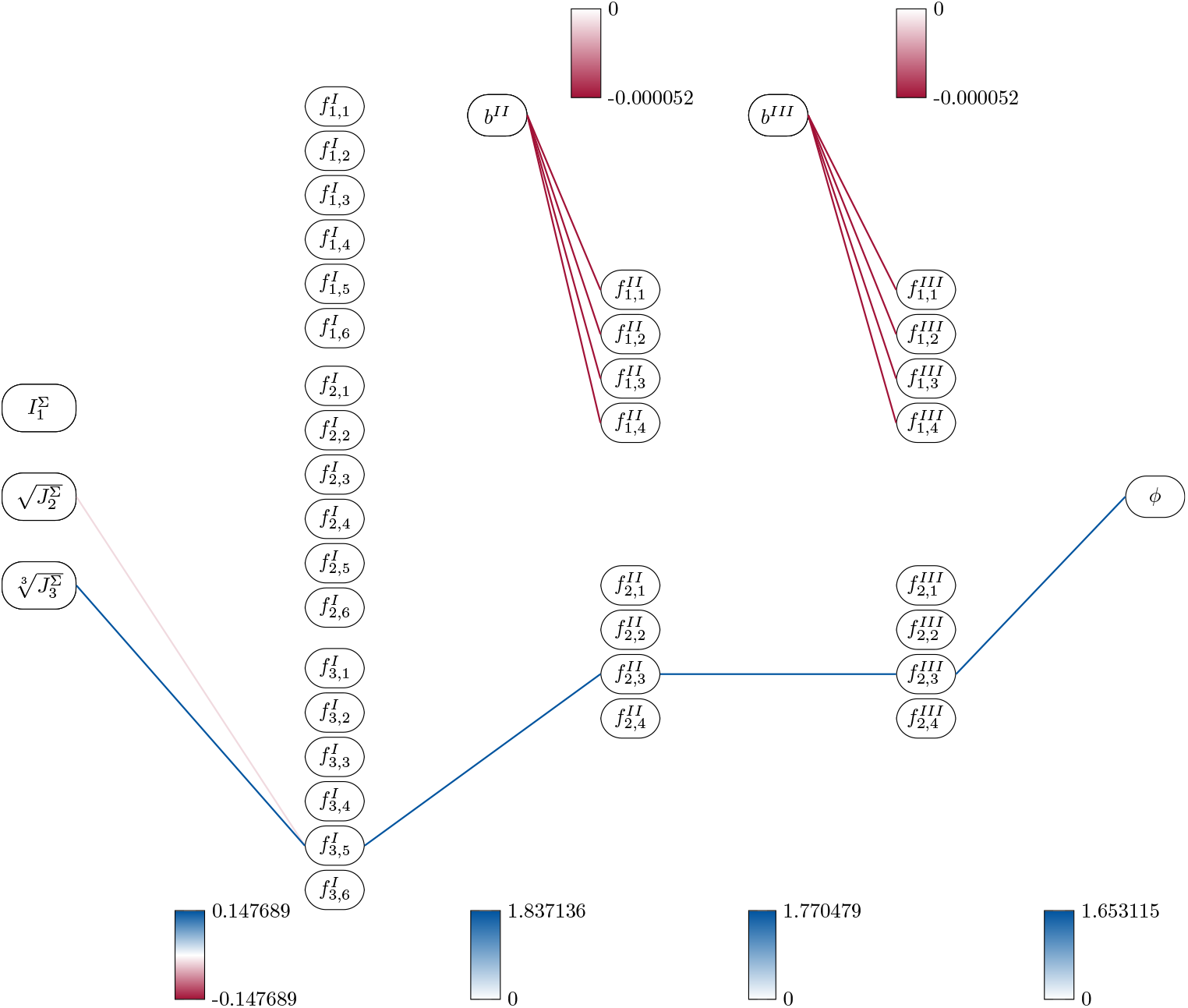
Model discovery - Trained dissipative network for silken tofu. The colors represent the magnitude of the final weights and biases, with blue indicating positive and red negative values, and white indicating that this path is inactive. The network discovers a single-term potential for silken tofu that only depends on the third deviatoric invariant 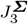 of the Mandel stress **Σ**.

For firm tofu in Figure 12, the dissipative potential is driven exclusively by the second deviatoric invariant 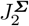,

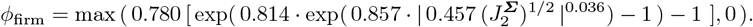

**Figure 12.**
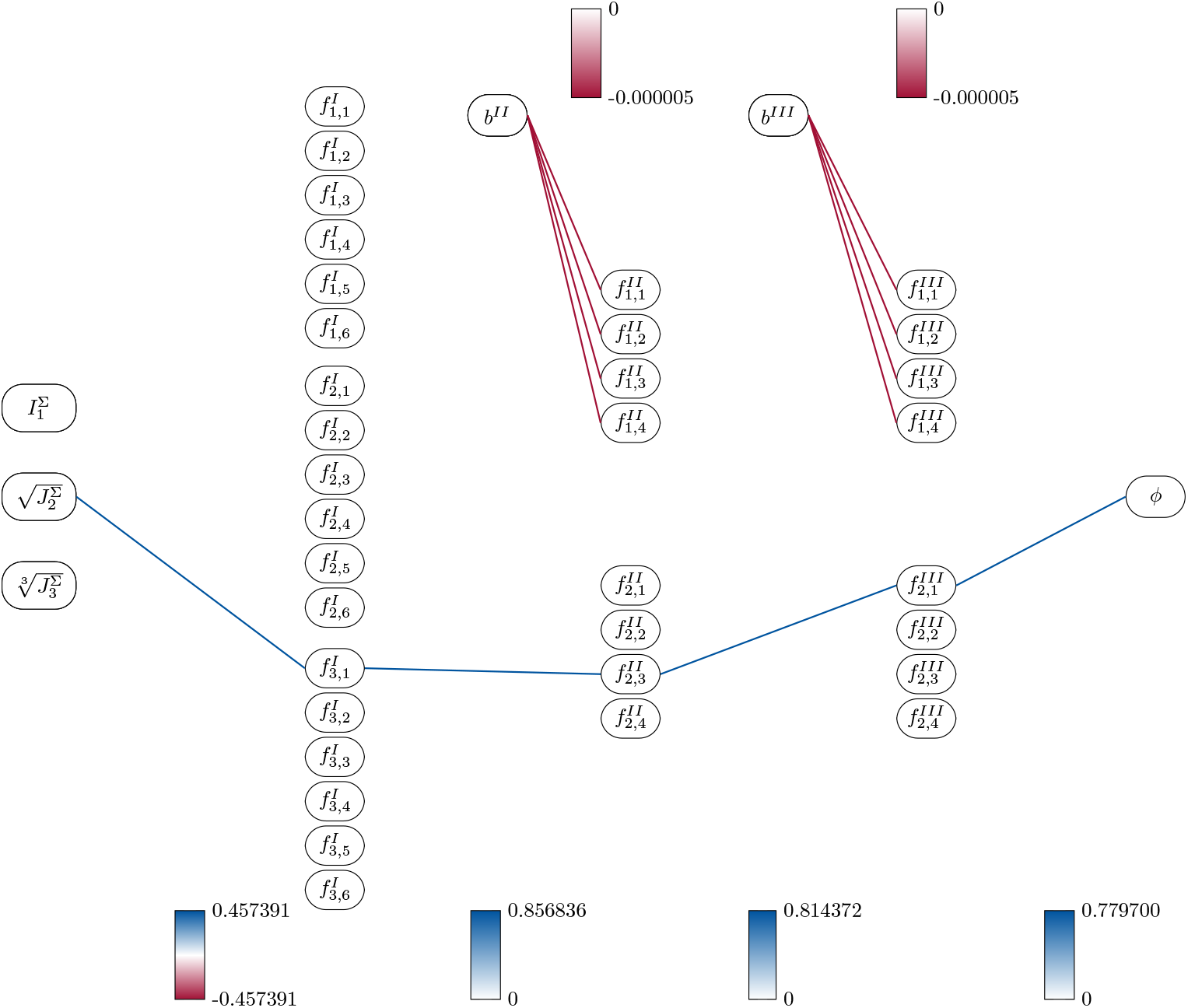
Model discovery - Trained dissipative network for firm tofu. The colors represent the magnitude of the final weights and biases, with blue indicating positive and red negative values, and white indicating that this path is inactive. The network discovers a single-term potential for firm tofu that depends on the second and third deviatoric invariants 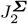and 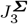of the Mandel stress **Σ**.

For extrafirm tofu in Figure 13, the dissipative potential is driven exclusively by the third deviatoric invariant 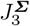,

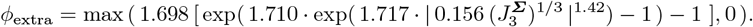

**Figure 13.**
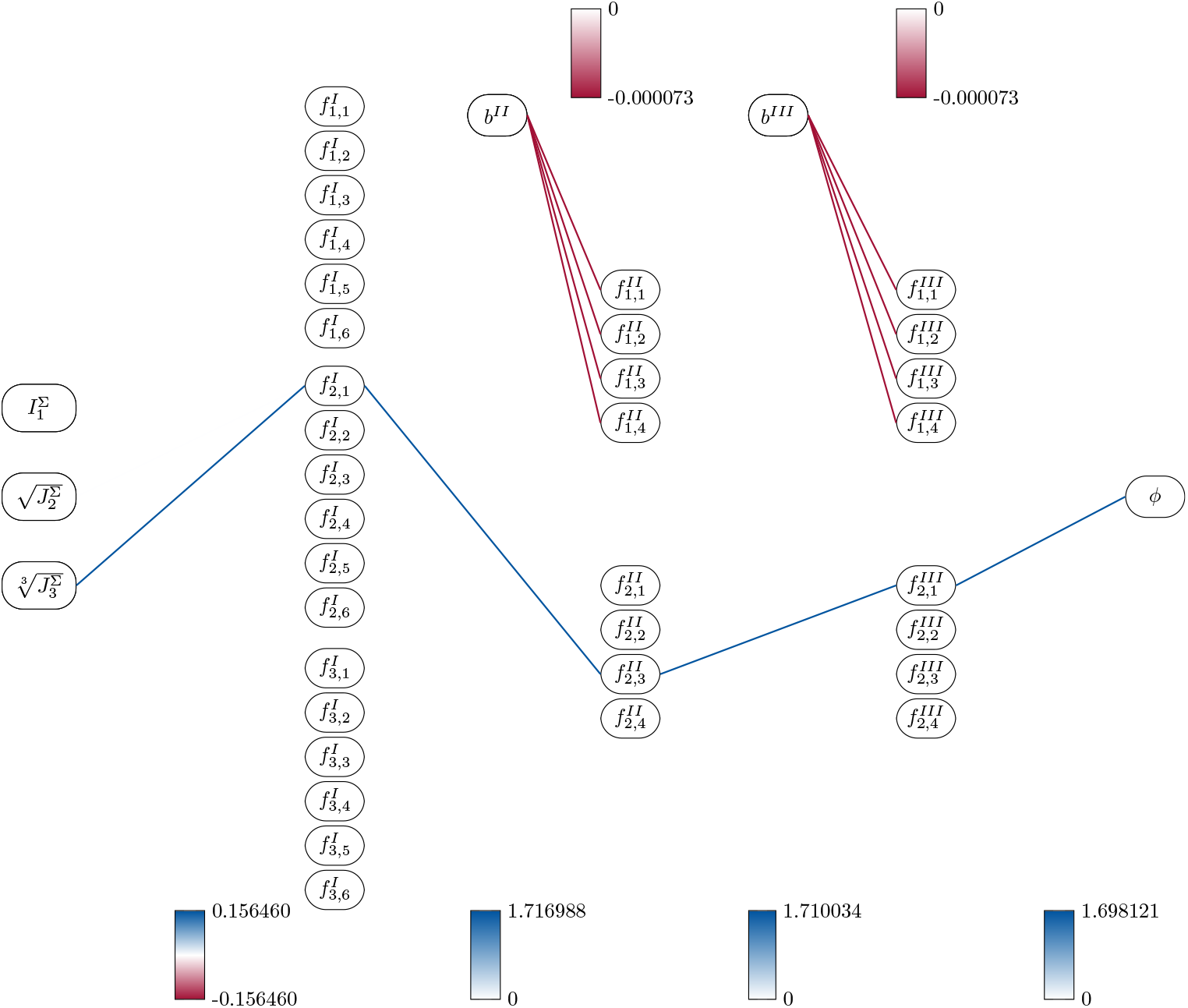
Model discovery - Trained dissipative network for extrafirm tofu. The colors represent the magnitude of the final weights and biases, with blue indicating positive and red negative values, and white indicating that this path is inactive. The network discovers a four-term potential for extrafirm tofu that depends on complex combinations of the second and third deviatoric invariants 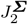 and 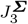 of the Mandel stress **Σ**.

Strikingly, for all three tofu types, the networks are sparse and discover the same functional form: a single term that contains weighted powers of the second and third deviatoric invariants 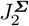 and 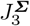 embedded in two nested exponential functions. None of the discovered potentials depends on the volumetric invariant 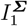, and none of them features any biases that would result in additive contributions.

Figure 14 illustrates the stresses and the inelastic deformation rates of the discovered models for silken, firm, and extrafirm tofu for a strain rate of 60%/s. The first row shows the applied stretches *λ*, followed by the total stress *S*, its equilibrium and non-equilibrium parts *S*^eq^ and *S*^neq^, and the inelastic deformation rate 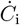. The color shades denote contributions from the three invariants 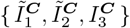 and 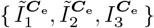 and 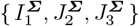. Strikingly, the predicted peak stress *S* in the second row increases notably with decreasing water content, from -3.90 kPa for silken to -19.62 kPa for firm and -33.85 kPa for extrafirm tofu, from left to right. The peak equilibrium stress *S*^eq^ in the third row increases also, from -1.26 kPa for silken to -2.44 kPa for firm and -2.76 kPa for extrafirm tofu, but less drastically than the total stress *S*. Notably, the peak non-equilibrium stress *S*^neq^ in the fourth row is almost entirely made up of second-invariant terms in 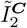 across all three tofu types. Similarly, the evolution of the inelastic right Cauchy tensor 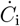 in the bottom row is made up only of second- and third-invariant terms in 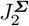and 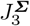, suggesting that the inelastic deformation remains entirely deviatoric. Importantly, as the water content decreases, from left to right, the equilibrium contribution *S*^eq^ to the total stress *S* decreases: It accounts for 1/3 of the total stress in silken, but for only 1/8 in firm and 1/12 in extrafirm tofu.

**Figure 14.**
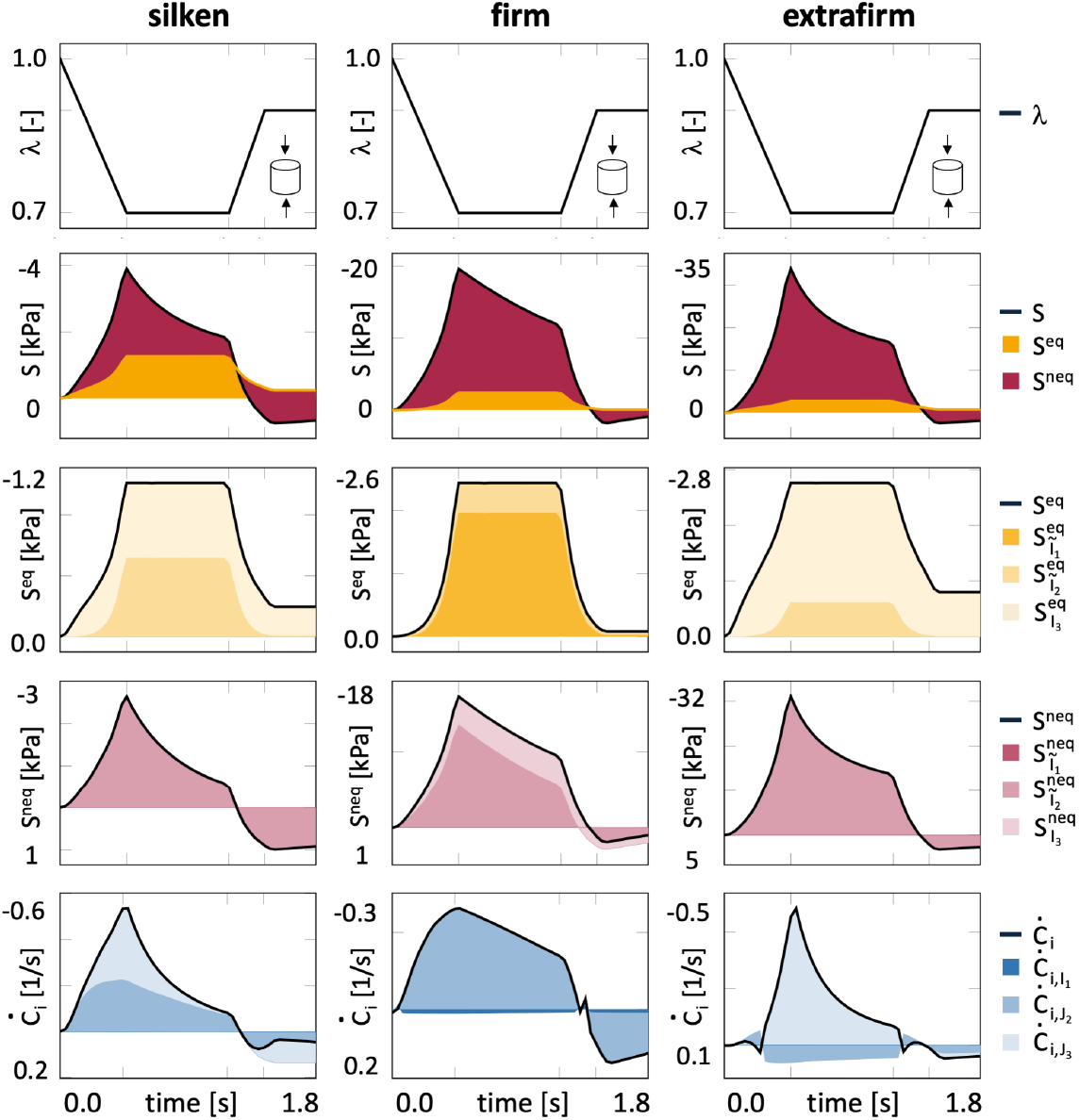
Model discovery - Predicted stresses and inelastic deformation rates of the three individual tofu models. Stress histories and evolution of the inelastic deformation rate for silken (left), firm (middle), and extrafirm (right) tofu for a strain rate of 60%/s. Applied stretches *λ* (top); total stress *S*, equilibrium stress *S*^eq^, and non-equilibrium stress *S*^neq^ (middle), and inelastic deformation rate 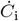 (bottom). Distinct patterns denote contributions from the three invariants 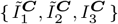 and 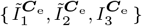 and 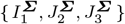.

### Discovering a single unified tofu model

Figure 15 summarizes the water-content dependency of the single unified tofu model. The rows correspond to the equilibrium, non-equilibrium, and dissipative potentials, the columns show the discovered dependencies of the individual terms, and the dashed lines indicate the water contents of our extrafirm, firm, and silken tofu. Interestingly, the majority of terms, six of eight, depend on the water content of tofu, while two terms do not: The equilibrium potential *ψ*^eq^ features water-content-independent quadratic first-invariant and logarithmic third-invariant terms, combined with water-content-dependent linear and quadratic second-invariant terms, where the first increases and the second decreases with water content. The non-equilibrium potential *ψ*^neq^ features water-content-dependent quadratic first-invariant and linear second-invariant terms, where both increase with water content. The dissipative potential *ϕ* features water-content-dependent exponential second- and third-invariant terms, where the first increases and the second decreases with water content.

**Figure 15.**
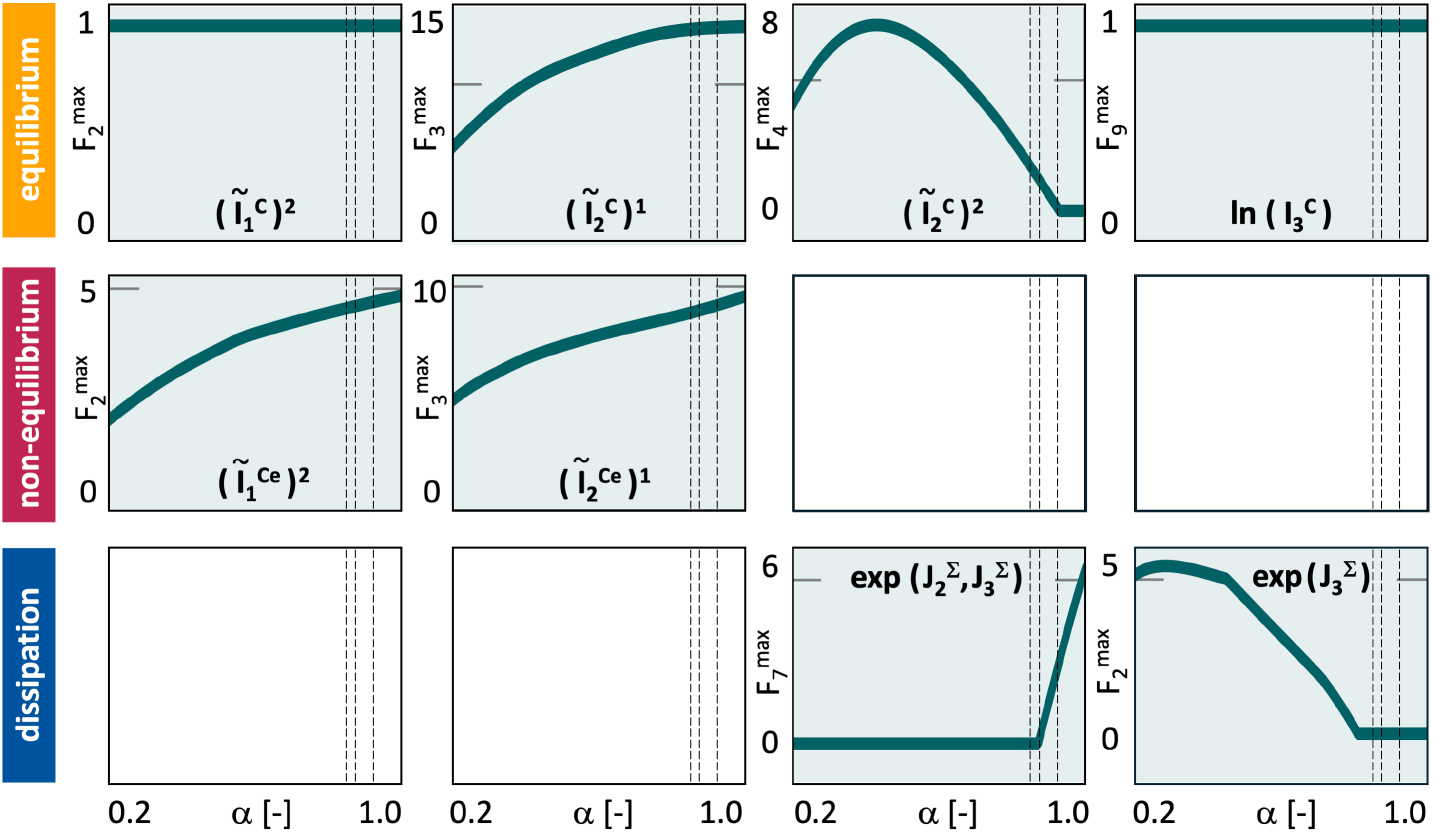
Model discovery - Water-content dependency of the single unified tofu model. Functional dependency of the equilibrium potential *ψ*^eq^ (top), non-equilibrium potential *ψ*^neq^ (middle), and dissipative potential *ϕ* (bottom) on the water content *α*. Columns show the discovered dependencies on the individual terms. Dashed lines highlight the water contents of extrafirm, firm, and silken tofu.

Figure 16 compares the computationally predicted stresses in green and the experimentally recorded stresses in gray. The rows correspond to silken, firm, and extrafirm tofu, and the columns represent distinct loading cases with varying stretch magnitudes and rates. In contrast to Figure 10, the stress predictions now result from a single model that includes the water content as an explicit input feature. This unified model results in a smaller coefficient of determination with R^2^ = 0.6900 compared to the single models with R^2^ = 0.7449, but still captures the underlying trends effectively. Appendix A provides additional details about the errors. Although the model is trained only on synthetic data, it reproduces the major experimental trends: For firm and extrafirm tofu, in the second and third rows, the computationally predicted stresses align closely with the experimentally recorded stresses across all strain rates, especially during loading and unloading. During holding, the predicted stresses follow the experimental averages, but display slightly less nonlinearity than the experimental measurements. For silken tofu, in the first row, the model overestimates the compressive peak stresses by a factor two, which is reflected in its negative R^2^-values. Despite this discrepancy, the silken tofu model captures the overall trends of nonlinear relaxation and decreasing stress magnitude with successive load cycles.

**Figure 16.**
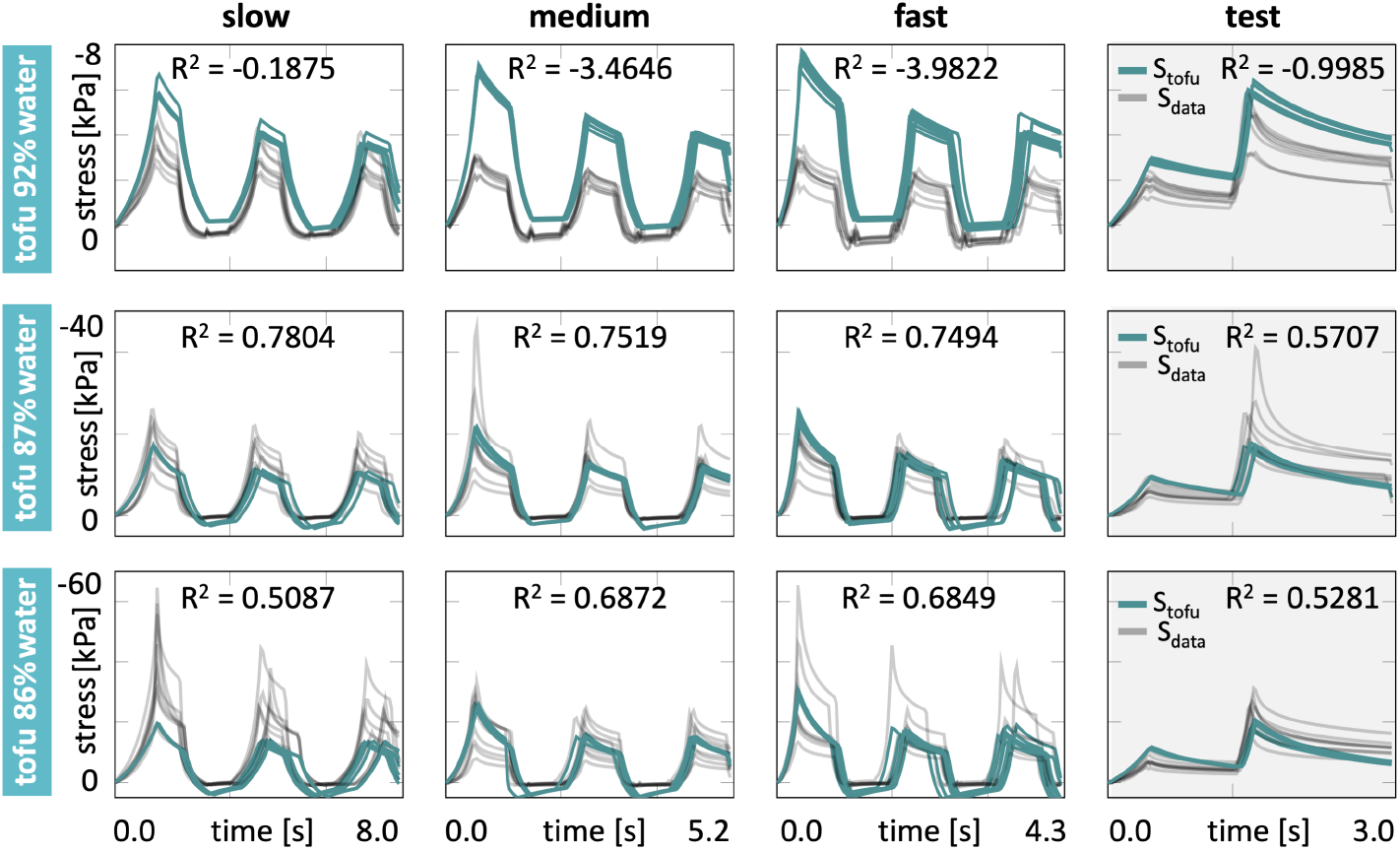
Model discovery - Experimentally recorded and computationally predicted stresses of the single unified tofu model. Stress histories *S*(*t*) for silken (top), firm (middle), and extrafirm (bottom) tofu. The first three columns show the triple-compression tests used as training data. The last column shows double-compression tests used as test data. Experimental data are plotted in gray, model predictions in green; R^2^ values indicate the goodness of fit.

## 6. Discussion

### Tofu is a nonlinear viscoelastic material

Tofu is an exceptional material made up of only soy beans and water. This simple yet tunable model system offers unique conditions to study composition–mechanics relationships. Our mechanical testing of three different tofu types across different water contents, strain magnitudes, and rates in Figures 8 and 9 reveals several interesting rheological characteristics: First, during the entirely loading path—loading, holding, and unloading–the response of tofu is markedly *nonlinear*. Second, during holding, stresses relax at constant stretch, and the response of tofu is notably *time-dependent*. During consecutive load cycles with similar peak stretches, the recorded peak stresses decrease, which supports the existence of stress relaxation. We conclude that the material behavior of tofu is made up of a history-independent *elastic* part and a history-dependent *inelastic* part. Our observations agree well with rheological tests of different foods throughout the past four decades [44, 64]. Specifically, our confined compression setup imposes non-permeable boundaries that prevent water from escaping the sample. This suggests that the observed stress relaxation arises primarily from the viscoelastic behavior of the solid [10]. rather than from the fluid-filled matrix [25]. We therefore first adopt the *theory of finite viscoelasticity* [65, 69] to explain the observed nonlinear time-dependent response.

### Tofu is loading-rate and water-content dependent

We further observe two general trends: stresses increase with increasing loading rate and with decreasing the water content. Figure 9 of the experimental data and Figure 10 of the discovered model highlight the loading-rate dependence from left to right and the water-content dependence from top to bottom. *The loading-rate* dependence agrees with our previous findings of fungi-based steak, where the stiffness increased by a factor five when varying the loading rate across four orders of magnitude [78]. This trend is visible in Figures 9 and 10, although less pronounced, since our loading rates only vary by a factor three. *The water-content dependence* is more obvious, for example, in the first column, where mean peak stresses are lowest in silken tofu with -3.74 ± 0.86 kPa, followed by firm tofu with -18.91 ± 5.50 kPa, and extrafirm tofu with -42.84 ± 16.22kPa. These stress ranges agree well with our previous tension, compression, and shear tests of firm and extrafirm tofu with mean stiffnesses of 26.4 ± 4.0 kPa and 27.5 ± 5.6 kPa [71] and with our rheologically measured storage moduli of 5.7 ± 0.5 kPa and 11.0 ± 1.0 kPa [22]. Our observations also agree with previous viscoelastic tests on lab-made tofu with different water contents [14], where the elastic, viscous, and relaxation parameters increased notably with decreasing water content [15]. We can potentially explain these observations with the *theory of finite poroelasticity* [19, 59], and propose to supplement the theory of finite viscoelasticity with volume fractions of the solid and fluid phases.

### Inelastic constitutive neural networks differentiate elastic and inelastic effects

Our experiments suggest that the constitutive behavior of tofu consists of elastic and inelastic parts. Yet, their individual contributions to the overall stresses remain poorly understood. To discover which fraction of the experimentally recorded stress is reversibly elastic and which is not, we utilize an *inelastic constitutive neural network* [1, 28] made up of feed forward networks for the equilibrium and non-equilibrium potentials combined with a recurrent neural network for the dissipative potential [29]. The consistent agreement between our computationally predicted and experimentally measured stretch–strain curves, across all three tofu types, for both training and test data in Figure 10, confirms that our network accurately captures the essential mechanical features of tofu. From the visualization of the equilibrium and non-equilibrium contributions to the overall stress in Figure 14, we conclude that the network autonomously differentiates elastic and inelastic effects. This differentiation provides an important physical insight: The equilibrium stress increases, from -1.26 kPa for silken to -2.44 kPa for firm and -2.76 kPa for extra firm tofu, as the water content decreases. This trend mirrors the classical constitutive theory for fluids, where a perfect fluid at rest cannot support shear stresses. In other words, its rheological model lacks a Hookean spring and only consists of a Maxwell element. Several rheological models with a single Maxwell model, a generalized Maxwell model, and a four-element Maxwell model have been proposed to explain the stress relaxation of tofu with different water contents [14]. Our automated discovery process captures these Maxwell-type characteristics [29], and naturally reproduces the gradual transition from a viscoelastic solid to a nearly fluid-like response.

### The elastic behavior of tofu is dominated by the second isochoric invariant

To characterize the isotropic elastic behavior of tofu, we have parameterized its equilibrium and non-equilibrium potentials in terms of two isochoric invariants and *one volumetric invariant* of the total and elastic right Cauchy Green tensors [33], 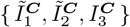 and 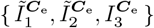. Yet, their individual contributions to the two elastic potentials are unknown. To discover which invariants drive the equilibrium and non-equilibrium responses, we consult our elastic networks in Figure 3 to identify the most relevant contributions out of 2^9^ = 512 possible combinations of terms [28, 29]. Despite quantitative differences, the discovered models for all three tofu types, silken, firm, and extrafirm, share several interesting trends: Across all tofu types, both elastic potentials, equilibrium and non-equilibrium, are dominated by the *second isochoric invariants*, 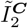 or 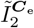, which measure the isochoric area change. The equilibrium potential *ψ*^eq^ features a quadratic second-invariant term for silken and extrafirm tofu, and a linear second-invariant term for firm tofu. The non-equilibrium potential *ψ*^neq^ features a linear second-invariant term for all three tofu types, and an additional quadratic second-invariant term for extrafirm tofu. The complete *absence of exponential terms*–in both potentials and across all three tofu types–is in line with our general understanding that exponential terms represent the strain-stiffening of elastin and collagen fibers in soft biological tissues [33], which are not present in tofu. The *dominance of the second invariant* [42] is in line with previously discovered models for human brain [48] and plant-based meat [50], and suggests that these hydrated soft materials share common rheological trends. Our results parallel classical constitutive modeling, where a single constitutive model can describe a family of materials simply by adjusting its parameter values. For example, in previous studies on lab-made tofu, a four-element Burgers model, a rheological model with two parallel dashpot-springs, was able to explain the rheological behavior of tofu with varying soymilk concentrations of {5%, 6%, 7%, 8%, 9% }, simply by varying the elastic and viscous parameter values and the relaxation time [15]. Our discovered models naturally reflect this principle: Tofu with varying solid volume fractions of {8%, 13%, 14% } results in varying parameterizations of the model, but not in fundamentally different functional forms.

### The dissipative behavior of tofu is driven by the second and third deviatoric invariants

To characterize the dissipative behavior of tofu, we have parameterized its dissipatve potentials in terms of *one volumetric invariant* and *two deviatoric invariants* of the Mandel stress [28], 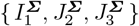 . Yet, their individual contributions to the dissipative potential are unknown. To discover which invariants drive the dissipative responses, we consult our dissipative network in Figure 4. Strikingly, across all three tofu types, the discovered dissipative potentials *ϕ* feature a single term that contains weighted powers of the second and third deviatoric invariants, 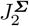 and 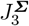, embedded in two nested exponential functions. None of the discovered potentials depends on the volumetric invariant 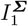, and none of them features any biases that would result in additive contributions. These observations confirm that our dissipative network autonomously eliminates irrelevant terms, and discovers sparse and interpretable models [56]: The first stress invariant 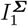 measures the hydrostatic stress, and its absence in the dissipative potential indicates a *pressure insensitivity*, unlike in the classical Drucker-Prager model [21]. The second stress invariant 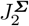 measures the magnitude of shear, and its presence in the dissipative potential indicates a *shear sensitivity*, like in the classical von Mises model [57]. The third stress invariant 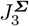 measures the mode of shear via the ratio between the principal stresses, and its presence in the dissipative potential indicates a *stress ratio sensitivity*, like in the classical Tresca [73] and Mohr-Coulomb models [16, 58]. While simple gels are often modeled as von-Mises type viscoelastic materials that depend exclusively on the second invariant, our dissipative network discovers a Lode-angle sensitive behavior [51] that depends on both the second and third invariants and distinguishes between tension-shear, pure shear, and compression-shear of the same shear magnitude [8]. These considerations underscore that our discovered models not only accurately reproduce the experimental behavior of tofu, but also offer interpretable, data-driven insight into its nonlinear viscoelastic behavior.

### The mechanics of tofu depend nonlinearly on its water content

Tofu is one of the simplest man-made foods; it consists of only two ingredients: soy beans and water. We can easily tune its consistency by adjusting its processing parameters: the higher the pressure and the longer its duration, the lower the water content [68]. Making and testing tofu with varying water content provides valuable insight into how we can modulate the rheology of food simply by adding or removing water [15]. From our three individually discovered tofu models, we conclude that the nonlinear and time-dependent behavior of tofu is highly sensitive to its water content. Yet, the explicit impact of water on the individual tofu models is unknown. A naive ad hoc explanation would be to consult the biphasic theory [59], adopt the concept of volume fractions [18], associate the equilibrium behavior with the solid and the non-equilibrium behavior with the fluid, and weigh both with their individual volume fractions [23], *S* = *ν*_s_ · *S*^eq^ + *ν*_f_ · *S*^neq^. In Table 1, the volume fractions of the fluid and solid are *ν*_f_ = { 0.92, 0.87, 0.86} and *ν*_s_ = 1 *− ν*_f_ = {0.08, 0.13, 0.14 }, for silken, firm, and extrafirm tofu. We can easily see that the *biphasic theory alone fails* to explain the observed stiffness variations: For example, the peak stresses in the first column of Figure 9, *S*_peak_ = *{−* 3.74 ± 0.86 MPa, *−* 18.91 ± 5.50 MPa, *−*42.84 ± 16.22 MPa } vary by more than an order of magnitude across the three tofu types, while the volume fractions vary by only six percent. This agrees with our previous observations of firm and extrafirm tofu for which the stiffness, storage, and loss moduli varied by a factor two, while the water contents differed by only one percent [22]. We conclude that volume-fraction-based scaling might be appropriate for linear mixtures with low-to-moderate water contents [19], but a more complex nonlinear water-content dependence is critical to accurately model highly hydrated soft materials like tofu.

### Feature network modeling reveals correlations between water content and mechanics

Feature networks learn non-linear mappings from raw input features to weights that parameterize or modulate terms of another network model. To explore how water content modulates the mechanics of tofu, we introduce a feature network that takes the scalar-valued water content *α* as input, and outputs the weights for each node of the three mechanical networks. Our results in Figure 15 confirm that the individual terms of the equilibrium, non-equilibrium, and dissipative potentials display different types of water-content dependency: Two terms are entirely independent of the water content, while four increase and two decrease with increasing water content. We conclude that incorporating the *water content as a feature* can successfully introduce material-specific variability into a unified constitutive model. The close agreement between the computationally predicted and experimental recorded stresses in Figure 16, especially for firm and extrafirm tofu, confirms that the feature network can indeed effectively modulate the mechanical response. Comparisons with other hydrated soft biological tissues suggest that neither a purely viscoelastic [11], nor a purely poroelastic model [25] can accurately capture this characteristic behavior, but that a poroviscoelastic model is needed instead [36]. Indeed, hydrated soft solids including the cornea, meniscus, intervertebral disk, articular cartilage, and hydrogels have long been known to display a complex nonlinear poroviscoelastic behavior [27]. Our feature network provides the inherent flexibility to model these complex rheologies. It naturally combines microstructural information and macrostructural experiments into a data driven, scalable approach that represents families of hydrated soft materials within a single feature-weighted model. More generally, our results suggest that feature network modeling is a flexible approach to integrate multimodal data and discover how various multiphysics inputs–water content, density, temperature, or fiber angles–modulate the mechanics and physics of soft matter systems.

### Limitations

We note a few limitations of our study: First, we have assumed that the *inelastic deformation* is isotropic and *volume preserving*, and that volumetric changes are associated with the elastic deformation only [33, 65, 69]. In a follow up study, we are currently performing both confined and unconfined compression tests on tofu to explore to which extent this assumption is justified. Second, our currently findings are restricted the *compressive loading* only. Although compression is the dominant loading mode during chewing and most relevant to our sensory perception of texture [77], we have previously tested firm and extrafirm tofu in compression and shear [22], and in tension, compression, and shear [71]; yet, testing silken tofu in tension remains challenging. Third, we have only tested over a *small range of loading rates*. Testing food at loading rates across several orders of magnitude and observing fluid flow can help delineate poroelastic and viscoelastic phenomena [77]. Fourth, we have assumed that water in tofu exists only in a single state [19], freely moving. In reality, tofu is a hydrated soy protein gel that contains water in three distinct states, *bound, immobilized*, and *free* [83], where bound and immobilized water modulate the elastic response, while free water modulates the inelastic response [52]. A next logical step would be to characterize the volume fractions of bound, immobilized, and free water [36, 37], then integrate all three volume fractions as distinct features of the network, and discover how each contribution modulates the behavior of tofu, and of hydrated soft solids in general. Finally–and this is probably more a philosophical question than a true limitation–in line with the literature [9, 24, 47], we have used the terminology *discover* in the context of discovering sparse, interpretable functional representations of potentials within a pre-specified constitutive ansatz. Following the common approach, we reverse-engineer this pre-defined ansatz from popular constitutive terms and functions and generalize it to a library of activation functions that satisfy thermodynamics and physical constraints by design [38, 40, 54].

## Conclusion

Tofu is a hydrated protein gel composed only of soybeans and water. It provides a minimal model system to explore how material composition governs the mechanics and physics of food. A series of compression tests of three tofu types reveals its strong nonlinearity, its pronounced stress relaxation, and its more than ten-fold increase in peak stress with only six percent decrease in water content. Using automated model discovery, we discover inelastic constitutive models and parameters that accurately explain both our training and test data of silken, firm, and extrafirm tofu across three loading magnitudes and five loading rates. The discovered models autonomously separate elastic and inelastic behavior and reveal that the elastic response is dominated by the second isochoric invariant, while the inelastic response is governed by a combination of the second and third deviatoric invariants. Water content modulates the magnitude of these mechanisms, but not their functional form. This motivates the integration of all results into a single water-content weighted model for different tofu types. Our results position tofu as a simple quantitative benchmark for nonlinear poroviscoelastic solids and demonstrate how physics-informed machine learning can expose the underlying constitutive structure of hydrated soft materials. Knowing the precise rheology of tofu is essential to reliably reproduce, engineer, and design the texture of alternative protein products in the benefit of human and planetary health. More broadly, our results suggest that combined feature–mechanics networks can uncover complex multiphysics behavior in hydrated soft materials, including biopolymer hydrogels, engineered protein networks, and hydrated porous composites.

**Figure B.17.**
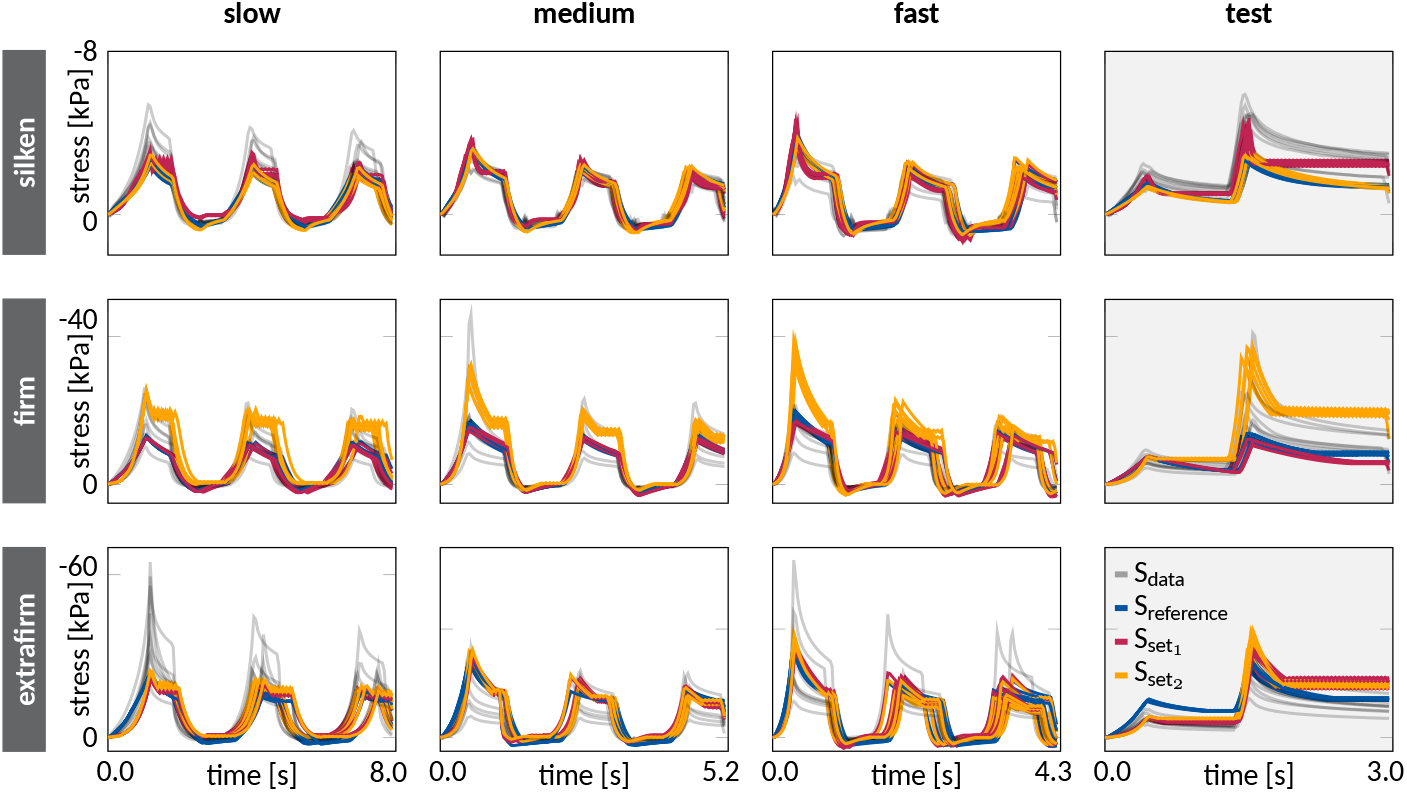
Preliminary sensitivity analysis for varying network architectures. Stress histories *S*(*t*) for silken (top), firm (middle), and extrafirm (bottom) tofu. Experimental data are plotted in gray; reference data from Figure 10 with {18, 8, 8} neurons per layer are plotted in blue; predictions with {30, 16, 12} and {4, 4, 2} neurons per layer are shown in red as set 1 and in orange as set 2.

## Appendix A. Errors of discovered individual and unified models

To quantify the performance of our discovered models, we calculate the error between the stress of the model *S* and the data Ŝ across all data points *n*_data_. In addition to the coefficients of determination R^2^ reported in Figures 10 and 16, we report the mean absolute error, 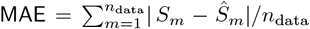, the root mean squared error, 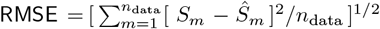, and normalized root mean squared error, NRMSE = RMSE*/*max { *Ŝ*}, where max { *Ŝ*} denotes the maximum experimental stress. Table A.2 summarizes the errors for the discovered individual tofu models in Figure 10 and for the discovered unified tofu model in Figure 16.

**Table A.2:** Errors of discovered models. Mean absolute error MAE, root mean squared error RMSE, and normalized root mean squared error NRMSE for the discovered individual tofu models in Figure 10, left, and for the discovered unified tofu model in Figure 16, right.

| individual | error | slow | medium | fast | test | unified | error | slow | medium | fast | test |
| --- | --- | --- | --- | --- | --- | --- | --- | --- | --- | --- | --- |
| silken | RMSE | 0.633 | 0.298 | 0.460 | 1.206 | silken | RMSE | 1.473 | 2.318 | 2.643 | 1.633 |
|  | MAE | 0.402 | 0.223 | 0.322 | 0.970 |  | MAE | 1.182 | 1.893 | 2.150 | 1.435 |
|  | NRMSE | 0.118 | 0.095 | 0.133 | 0.205 |  | NRMSE | 0.274 | 0.742 | 0.762 | 0.277 |
| firm | RMSE | 2.911 | 2.891 | 2.476 | 4.203 | firm | RMSE | 2.867 | 3.140 | 3.056 | 3.880 |
|  | MAE | 1.871 | 1.684 | 1.625 | 2.707 |  | MAE | 2.131 | 2.231 | 2.321 | 2.728 |
|  | NRMSE | 0.112 | 0.063 | 0.113 | 0.103 |  | NRMSE | 0.111 | 0.068 | 0.141 | 0.095 |
| extrafirm | RMSE | 5.746 | 4.877 | 5.200 | 4.644 | extrafirm | RMSE | 7.632 | 4.030 | 5.405 | 4.080 |
|  | MAE | 3.400 | 3.386 | 3.659 | 3.922 |  | MAE | 4.982 | 3.290 | 3.642 | 3.297 |
|  | NRMSE | 0.089 | 0.147 | 0.080 | 0.148 |  | NRMSE | 0.118 | 0.122 | 0.083 | 0.131 |

## Appendix B. Preliminary sensitivity analyses

Prior to generating the main results of this study, we performed a systematic series of sensitivity analyses. Here we showcase three exemplary studies that, amongst many others, have guided our selection of network architectures, activation functions, and hyperparameters.

**Figure B.18.**
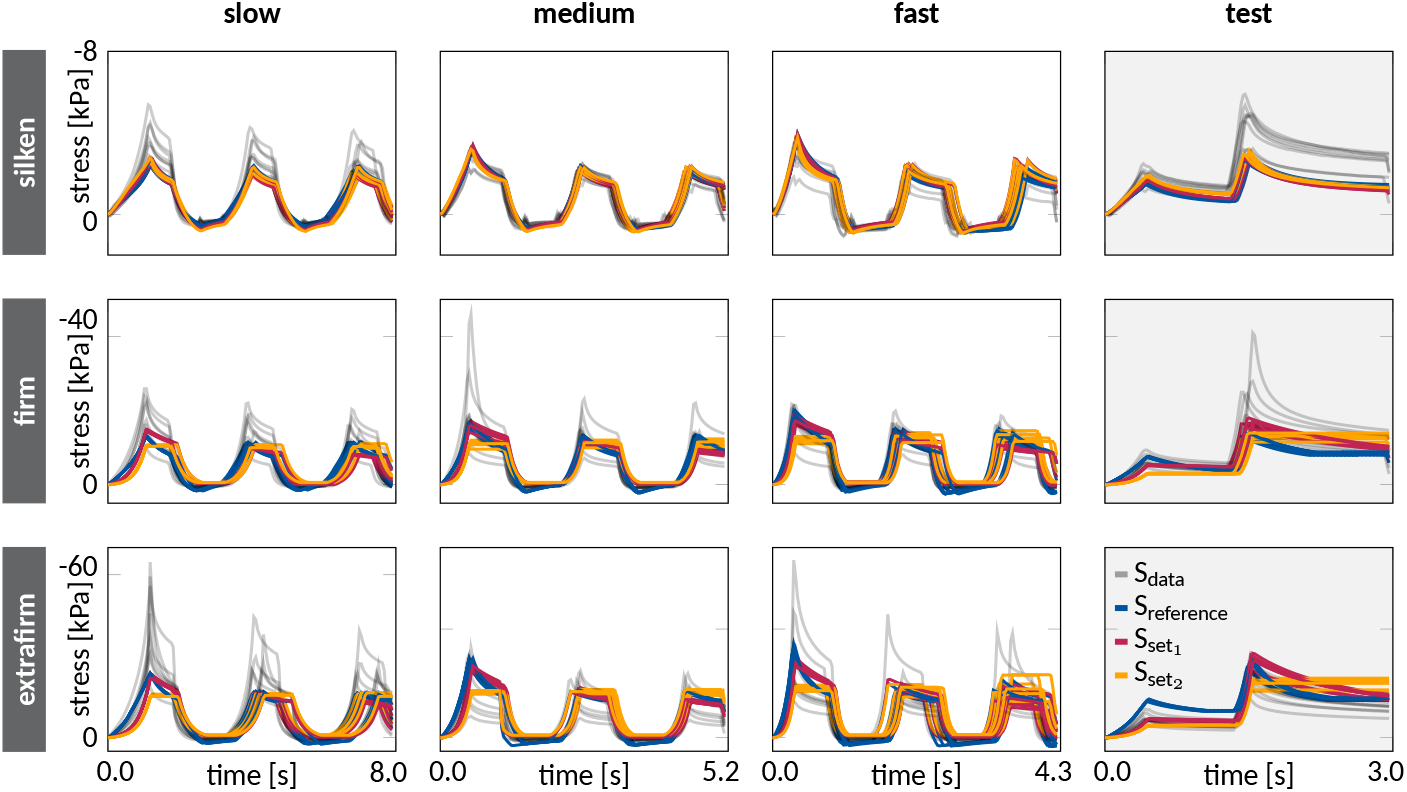
Preliminary sensitivity analysis for varying activation functions. Stress histories *S*(*t*) for silken (top), firm (middle), and extrafirm (bottom) tofu. Experimental data are plotted in gray; reference data from Figure 10 with trainable parameter *p* are plotted in blue; predictions with fixed parameters *p* = 1 and *p* = 2 are shown in red as set 1 and in orange as set 2.

**Figure B.19.**
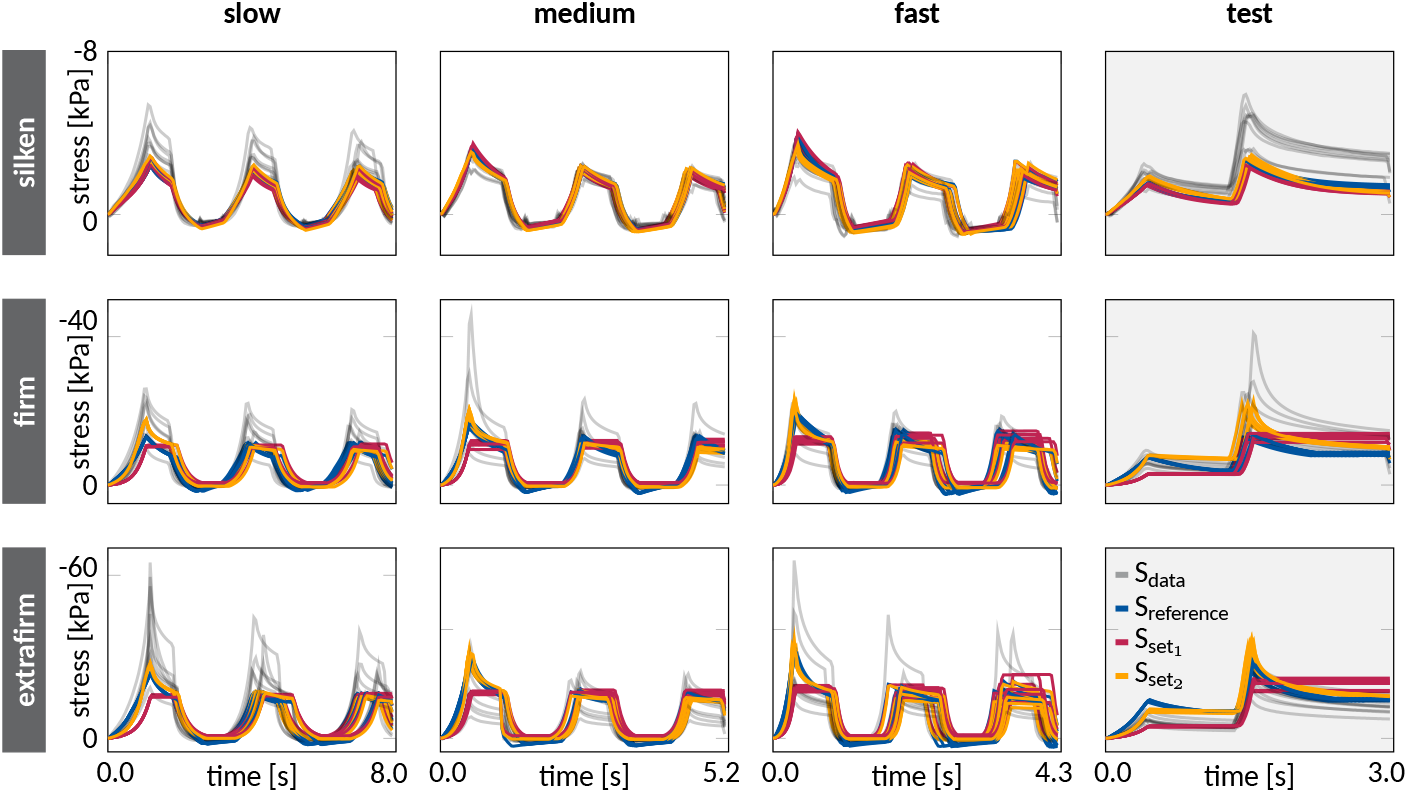
Preliminary sensitivity analysis for varying regularization parameters. Stress histories *S*(*t*) for silken (top), firm (middle), and extrafirm (bottom) tofu. Experimental data are plotted in gray; reference data from Figure 10 with regularization parameters *α*_1_ = *α*_2_ = 0.0001 are plotted in blue; predictions with *α*_1_ = *α*_2_ = 0.001 and *α*_1_ = *α*_2_ = 0.00001 are shown in red as set 1 and in orange as set 2.

### Network architectures

Figure B.17 summarizes our preliminary sensitivity analysis for varying network architectures. Specifically, we study a finer network with {30, 16, 12} neurons per layer and a coarser network with {4, 4, 2} neurons per layer, compared to our reference network with {18, 8, 8} neurons per layer that achieved a goodness of fit of R^2^ = 0.7449. The finer {30, 16, 12} network results in a reduced goodness of fit of R^2^ = 0.7085, associated with the stress predictions in red. The coarser {4, 4, 2} network results in a similarly reduced goodness of fit of R^2^ = 0.7091, associated with the stress predictions in orange. Interestingly, both network architectures lead to numerical instabilities, most visible in the results for firm and extrafirm tofu. These instabilities could be a result of time-step sensitivity related to the explicit time integration and might require a reduced time step size or more steps in the staggered training approach.

### Activation functions

Figure B.18 summarizes our preliminary sensitivity analysis for varying activation functions. Specifically, we vary the activation functions in the first layer of the dissipative network by fixing the powers to *p* = 1 and *p* = 2, instead of leaving *p* as a trainable parameter which achieved a goodness of fit of R^2^ = 0.7449. Fixing the powers to *p* = 1 and *p* = 2 reveals similar trends and a notable reduction of the goodness of fit to R^2^ = 0.7212 for *p* = 1, associated with the red curves, and even further to R^2^ = 0.6586 for *p* = 2, associated with the orange curves. For the latter, non-linear effects almost vanish entirely for firm and extrafirm tofu. Overall, our selection of activation functions is guided by thermodynamical considerations, and is in line with previous elastic [47, 48, 49] and dissipative [28, 30, 31] networks similar to Figures 3 and 4 that have demonstrated high robustness and reproducibility.

### Regularization parameters

Figure B.19 summarizes our preliminary sensitivity analysis for varying regularization coefficients *α*_1_ and *α*_2_ that fine-tune the level of *L*_1_ and *L*_2_ regularization. Specifically, we set both hyperparameters a magnitude higher and lower than our selected reference values of *α*_1_ = *α*_2_ = 0.0001 that achieved a goodness of fit of R^2^ = 0.7449 Higher regularization coefficients of *α*_1_ = *α*_2_ = 0.001 result in a reduced goodness of fit of R^2^ = 0.6556 with stress predictions that display almost no nonlinearity for firm and extrafirm tofu, shown in red. Lower regularization coefficients of *α*_1_ = *α*_2_ = 0.00001 result in a similar goodness of fit of R^2^ = 0.7510 as the reference model, with stress predictions, shown in orange, that predict similar features as the reference model, shown in blue.

Taken together, these results illustrate the sensitivity of network training with respect to the network architecture, activation functions, and regularization parameters. While we only show a few selected cases here, we have performed a wide variety of preliminary studies and recommend systematic sensitivity analyses before training the final model.

## Acknowledgements

We thank Sofie Danowski for assisting with the hyperparameter optimization as part to this study. This work was supported by the DFG Project SI 1959/12-1 466117814, by the DFG TRR 280 417002380, by the publication prize of the University of Wuppertal, by the Stanford Bio-X Snack Grant 2025, by the NSF CMMI Award 2320933, and by the ERC Advanced Grant 101141626.

## CRediT authorship contribution statement

BB: Conceptualization, Methodology, Software, Formal analysis, Data Curation, Investigation, Validation, Writing Original Draft, Writing Review and Editing. JWS: Funding acquisition, Writing Review and Editing. HH: Conceptualization, Methodology, Software, Formal analysis, Validation, Writing Review and Editing. EK Conceptualization, Resources, Writing Review and Editing.

## Data availability

Our source code and examples are available at https://doi.org/10.5281/zenodo.16993236.

